# Prefrontal–accumbens neural dynamics abnormalities in mice vulnerable to develop food addiction

**DOI:** 10.1101/2022.11.22.517466

**Authors:** Pablo Calvé, Thomas Gener, Laura Ribalta-Vilella, Sami Kummer, Pau Nebot, Elena Martín-García, M. Victoria Puig, Rafael Maldonado

## Abstract

Food addiction is characterized by a loss of behavioral control over food intake and is closely associated with several eating disorders, including obesity and binge eating. Despite its high prevalence, the underlying neural mechanisms of food addiction are still unresolved. We trained mice in an operant paradigm for 110 days to promote the development of food addiction. Then, we classified mice as addicted and extreme non-addicted based on three addiction criteria and recorded neural activities in the prelimbic medial prefrontal cortex (mPFC) and nucleus accumbens (NAc) core through electrophysiology *in vivo*. Addicted mice presented disrupted mPFC-to-NAc signaling at high frequencies (hfo 150-200 Hz) during decision-making to obtain food. Moreover, addicted mice exhibited reduced low gamma oscillations and theta-gamma coupling in the NAc during reward expectancy. Disrupted mPFC-to-NAc connectivity and gamma synchrony in the NAc correlated with increased reinforcement levels, unraveling the functional relevance of these alterations. The cannabinoid type-1 (CB1) receptor antagonist rimonabant rescued neural alterations observed in the addicted mice.Reinforcement levels were reduced after rimonabant administration and the directionality of signals and oscillatory activity in the NAc were reversed in addicted mice during decision-making and reward expectation, respectively. These findings suggest that disrupted mPFC-NAc neural dynamics are candidate mechanisms underlying specific behavioral alterations associated with food addiction. The elucidation of these novel communication mechanisms between the mPFC and the NAc will provide advances towards future development of new therapeutic interventions for food addiction and related disorders.

## Introduction

Food addiction is a disorder characterized by loss of behavioral control and compulsive food intake that is becoming increasingly prevalent in modern societies resulting in elevated health and socioeconomic costs^1,2^. It has been proposed that certain types of food, particularly highly palatable and fattening foods, could be factors that trigger addiction in vulnerable individuals, which may lead to overeating and obesity^3,4^. The term food addiction is still controversial and has not been included in the 5^th^ edition of the Diagnostic and Statistical Manual of Mental Disorders (DSM-5)^5^, although the Yale Food Addiction Scale v2.0 (YFAS v2.0) has been developed to precisely diagnose this disorder^6^. The concept of food addiction has been discussed in multiple scientific reviews^7–9^, with a recent particular interest on the link between food and drug addiction^10^. Indeed, both addictive processes could share common genetic and neurobiological mechanisms with overlapping dysregulations in the reward circuit, hijacking neurobiological pathways associated with reward sensitivity, incentive motivation, and inhibitory control, among others^5,9,11–14^.

Addiction is a complex multifactorial disorder with the participation of multiple environmental and genetic factors^15^. The behavioral abnormalities underlying this disorder seem to be mediated by key neurotransmitters that influence rewarding circuits and favor the transition to addictive behaviors. These processes induce pathological dysregulation of interconnected cortico-limbic areas, where the prelimbic medial prefrontal cortex (mPFC) and the nucleus accumbens (NAc) core play relevant roles^16,17^. The mPFC is a key structure that integrates reward-seeking and inhibitory control processes^18–20^. Previous studies have revealed a hypoactivation of the mPFC in addicted humans^17^ and animals with a compulsive cocaine self-administration^21^. The NAc is a crucial hub of communications involved in emotional and rewarding processing. Indeed, imaging studies in humans have demonstrated an association between NAc activity and subjective reward descriptors^22–24^. Interestingly, the mPFC and the NAc are densely interconnected through specific projections that play crucial roles in the inhibitory control of behavior. The mPFC sends glutamatergic efferent projections to the NAc while the NAc also reaches the cortex through glutamatergic projections, and addiction-related behaviors have been linked to abnormal neural activities of this circuit^18,25,26^. However, the role of this possible disrupted mPFC-NAc neural dynamics in food addiction has not been yet investigated.

Diverse innovative scientific tools have been employed to investigate specific neural circuits involved in food addiction^15^. Our previous studies have demonstrated the involvement of the mPFC-to-NAc pathway in food addiction, revealing that the inhibition of glutamatergic projections underlies vulnerability to develop food addiction in mice^27^. In addition, we have recently identified specific mechanisms of mPFC-dependent behavioral control in food addiction development in mice and humans^28^. Therefore, the communication between the mPFC and the NAc seems crucial in food addictive-like behaviors^15,27^. In agreement, neuronal spiking activity in the mPFC and NAc fluctuates between anticipatory and post-reward responses during drug self–administration regimes in animals^29–32^. Recent studies have focused on understanding oscillatory activities between the mPFC and NAc during drug rewarding outcomes, suggesting that addictive-like behaviors may be associated with irregular theta and gamma oscillations^33^. However, circuit communication measures between the mPFC and NAc during food addiction remain unknown.

Both the mPFC and NAc are extensively modulated by the endocannabinoid system, which seems involved in the development of food addiction^34^. Thus, the cannabinoid type-1 (CB1) receptor is crucial for addictive-like behaviors and represents an interesting therapeutic target for food addiction^35^. Indeed, deletion of CB1 receptor activity prevented the transition from controlled to compulsive seeking of palatable food in mice^36^, specifically in mPFC glutamatergic neurons^27^. However, the functional consequence of CB1 receptor blockade in mPFC-NAc neural dynamics during food addiction has not been yet investigated.

The present study aimed to investigate mPFC-NAc circuit disturbances in mice vulnerable to develop food addiction and the rescuing abilities of CB1 blockade. For this purpose, mice were trained in an operant model that mimics behavioral anomalies associated with food addiction^36^, and neural activities were recorded in the mPFC and NAc in mice vulnerable or resilient to develop this addictive-like behavior.

## Methods

### Animals

Experiments were performed in C57BL/6J male mice (n=31). Mice (2-10 months old) were single housed under a 12h reverse dark/light cycle. Details are provided in the Supplementary Information.

### Surgeries and electrode implantation (Detailed in the *SI*)

Tungsten electrodes were unilaterally implanted in the prelimbic mPFC (AP: 1.98mm, ML: 0.3mm, DV: −2.30mm) and in the NAc core (AP:1.10mm, ML:1.0mm, DV:-4.60mm). Implants were fixed to the skull with micro-screws. At the end of the experiment, electrode placements were confirmed histologically.

### In vivo electrophysiological recordings (Detailed in the *SI*)

Electrophysiological recordings were conducted in the dark illumination cycle. Recordings were carried out with the multi-channel Open Ephys system at 0.1-6000Hz and a sampling rate of 30kHz. To obtain LFPs, electrophysiological recordings were sampled to 1kHz, decoded, and filtered. The frequency bands considered for specific band analyses were delta, theta, beta, low gamma (lgamma), high gamma (hgamma) and high frequency oscillations (hfo). MUA was estimated by subtracting the raw signal from each electrode with the signal from a nearby referencing electrode to remove artifacts related to the animal’s movement. Then, continuous signals were filtered between 450-6000Hz.

### Experimental design (Detailed in the *SI*)

C57BL/6J mice were trained in operant boxes for 110 days to develop food addictive-like behaviors^27,36^. Persistence of response, motivation and compulsivity were evaluated in the early and late periods. According to the late period, mice were classified and ordered on a gradual addiction scale that lead to extreme phenotypes: addicted and non-addicted mice. After recovery, recordings were performed in operant boxes adapted for electrophysiology (Fig. 1A-B, Supplementary Fig.1A). Details are provided in the Supplementary Information.

**Fig. 1.**
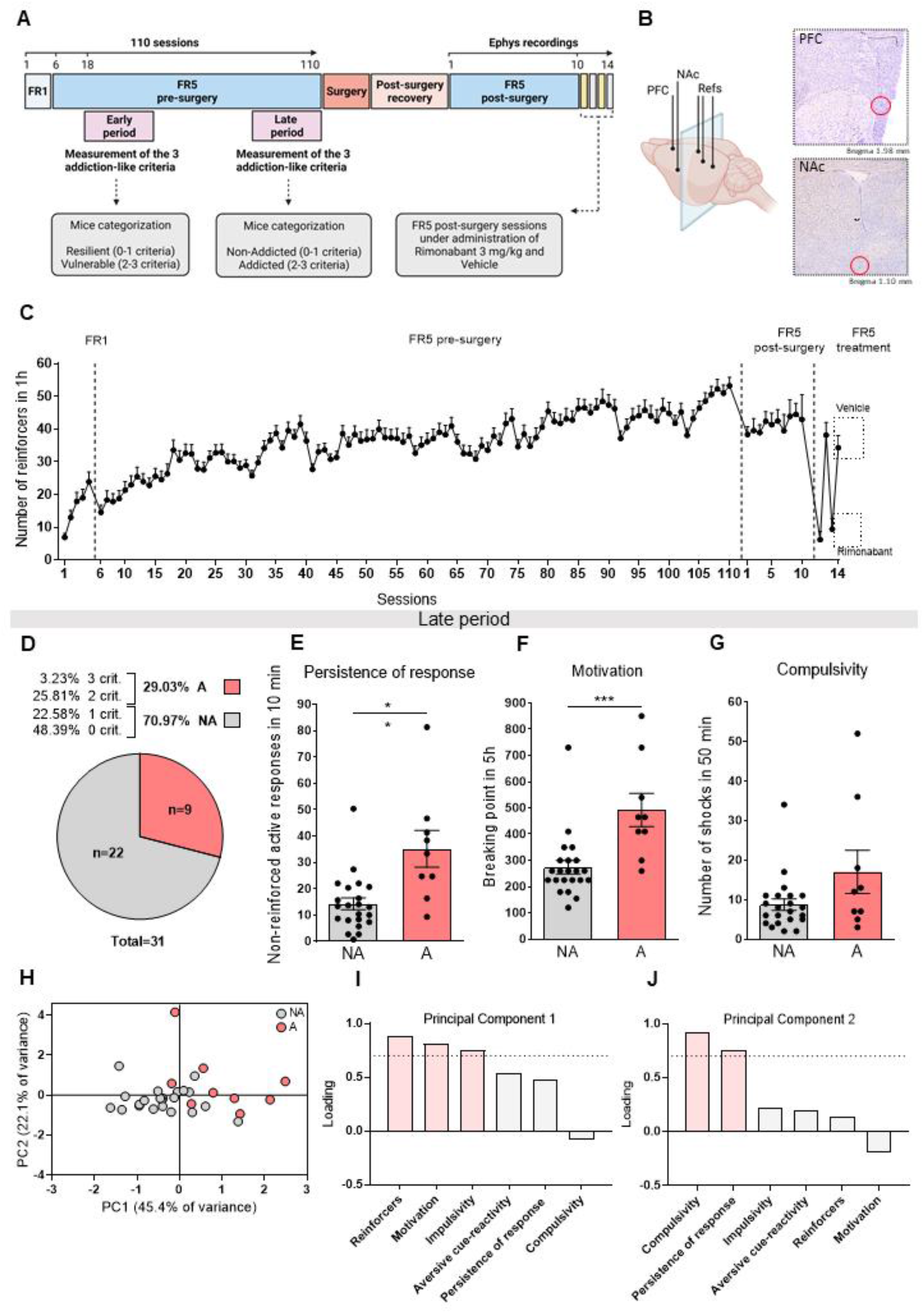
Identification of extreme phenotypes of addicted and non-addicted mice by long-term operant training. **A** Experimental design. Experimental sequence of the food addiction mouse model, electrophysiological recordings and pharmacological treatment. **B** Representative scheme and histological validation of electrode implantation in the mPFC and in the NAc. **C** Acquisition of operant training maintained by chocolate-flavored pellets. Mean number of reinforcers during the acquisition training in FR1 and FR5 schedule of reinforcement (n=31), during electrophysiological recordings (n=9-13), and during rimonabant treatment (n=9-11). Differences are reported as mean ±SEM. **D** Distribution of animals with different scores for addiction-like behavior according to late period. Animals were assigned to a criteria subgroup based on the amount of criteria met for which they scored equal or above the 75^th^ percentile. **E-G** A mice increased response in the three addiction-like criteria tests. **E** Persistence of response. **F** Motivation. **G** Compulsivity. Differences are reported as mean ±SEM, U Mann-Whitney, ** P<0.01, ***P<0.001. **H** Mice subjects clustered by presence of A or NA on the space yielded by two components of the PCA that account for the maximum data variance with factor loadings of principal component (PC) 1 (45.4%) and principal PC2 (22.1%). **I-J** The order of factor loading of the different variables in the (**I**) PC1 and (**J**) PC2 is represented. The dashed horizontal line marked loadings >0.7, mainly contributing to the component.

## Results

### Identification of extreme phenotypes of addicted and non-addicted mice by long-term operant training

We trained a cohort of 31 mice in a long-operant behavioral model of food addiction^27^ (Fig.1A). After the training, animals were surgically implanted with electrodes into the NAc and mPFC (Fig.1B, Supplementary Fig.1B). As expected, pellet intake progressively increased across sessions in all the mice (Fig.1C). No differences were detected in addiction-like criteria between mice during the early period (Supplementary Fig.1B-D). By contrast, at the late period around thirty percent (29.03%) of mice reached two or three addiction-like criteria and were considered as addicted. The remaining animals (70.97%) reached none or one criterion and were considered non-addicted (Fig.1D). Consistently, addicted mice obtained, in the late period, more reinforcers and performed more lever-presses than their non-addicted peers (Supplementary Fig.1E-F). To further characterize addictive-like behaviors, we investigated other relevant phenotypic traits beyond the three main addiction criteria, including impulsivity and aversive cue-reactivity. Addicted mice also showed significant differences in phenotypic traits (impulsivity and aversive cue-reactivity) associated with the development of food addiction during the late period. Addicted mice showed increased impulsivity (Supplementary Fig.1G) and non-significant decreased aversive cue-reactivity as revealed by enhanced food-seeking (Supplementary Fig.1H-I). Body weight increased similarly in both vulnerable and resilient mice (Supplementary Fig. 1J-K), indicating that food addictive-like behavior was independent of body weight changes. To gain deeper insight into the main factors contributing to the development of food addiction, we performed principal component analysis (PCA) of the different behavioral responses. PCA allowed classifying the mice into vulnerable and resilient phenotypes to develop food addiction combining the three addiction criteria and the two associated phenotypic traits (Fig. 1H). The percentage of variance explained by the two principal components was 45.4% (PC1) and 22.1% (PC2). The main variables contributing to PC1 were number of reinforcers, motivation criterion, and impulsivity trait, whereas compulsivity and persistence of response criteria were the predominant variable in PC2 (Fig.1I-J). As the number of reinforcers was the variable with the highest score of the PC1 (0.88) (Fig.1I), we focused the electrophysiological analysis on the reinforcement sessions during FR5.

### Prefrontal-accumbens synchrony at gamma and high frequencies are disrupted in addicted mice during decision-making and reward expectation

We ordered the mice in a quantitative gradual addiction scale (Fig.2A).This scale is calculated according to a specific formula (Supplementary information). Thus, we selected mice at the extremes with the lowest (extreme non-addicted mice) and the highest (addicted mice) scores to investigate the neurophysiological signatures underlying food addiction. Extreme non-addicted and addicted mice (Fig.2A) underwent stereotaxic surgery for electrode implantation in the mPFC and the NAc (Fig.1A,B).

**Fig. 2.**
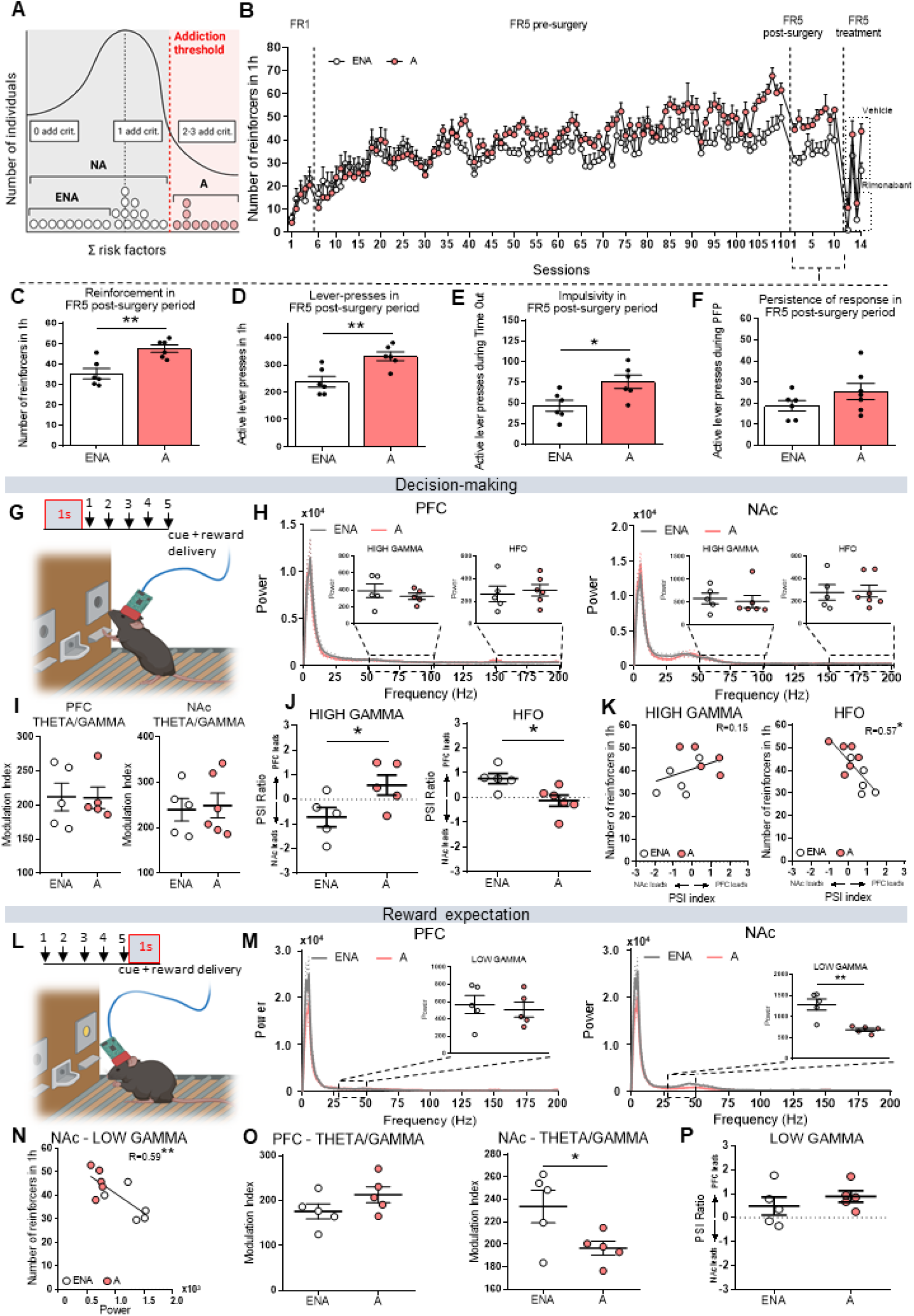
Prefrontal-accumbens synchrony at gamma and high frequencies are disrupted in addicted mice during decision-making and reward expectation. **A** Representative scheme of the selection of animals situated at the extremes of the curve, according to addictive-like criterions and phenotypic traits. Out of 31 animals, 22 were categorized as NA (0 and 1 criteria accomplished) and 9 were categorized as A (2 and 3 criteria accomplished). 9 extreme non-addicted (ENA) that accomplished 0 addiction-like criteria and obtained a low score in the addiction classification were selected in parallel with 9 A animals for electrode implantation. **B** Acquisition of operant training maintained by chocolate-flavored pellets in ENA (n=9) and A (n=9) mice. Mean number of reinforcers during the acquisition training in FR1 and FR5 schedule of reinforcement, during electrophysiological recordings, and during rimonabant treatment. Differences are reported as mean ±SEM. **C-F** During recordings of the FR5 post-surgery period, A mice maintained the addictive-like behavior during the operant sessions showing an increased **C** reinforcement, **D** number of lever-presses, and **E** impulsivity, in comparison with ENA mice (unpaired t-test, *P<0.05, **P<0.01). **F** Persistence of response was not significantly different between A and ENA mice. **G** Schematic diagram illustrating decision-making epochs. Decision-making epochs were defined as the second prior to the first of the five lever-presses in reward trials (5 lever-presses + cue light and reward delivery). **H** Power spectra of neural signals in the mPFC and the NAc in ENA and A mice during decision-making. Power values amplification of hgamma and hfo is also shown. **I** Quantification of local theta-gamma cross-frequency coupling in the mPFC and the NAc during decision-making. **J** PFC-NAc circuit communication (PSI) at hgamma and hfo frequencies during decision-making (unpaired t-test, *P <0.05). **K** Pearson correlations between PFC-NAc circuit communication at hgamma and hfo; and the number of reinforcers acquired in the FR5 operant sessions. mPFC to NAc hfo PSI that emerged during decision making correlated negatively with reinforcement levels in ENA and A mice together (Pearson correlation, *P< 0.05). **L** Schematic diagram illustrating reward expectation epochs. Reward expectation epochs were defined as the second occurring immediately after the fifth lever press, when the cue light turns on and reward is delivered. **M** Power spectra of neural signals in the mPFC and the NAc in ENA and A mice during expectation of reward. Power values amplification of lgamma is also shown (unpaired t-test, **P<0.01). **N** Pearson correlation between NAc lgamma power and the number of reinforcers acquired in the FR5 post-surgery operant sessions. Lgamma power negatively correlated with reinforcement levels in ENA and A mice together (**P<0.01). **O** Quantification of local theta-gamma cross-frequency coupling in the mPFC and the NAc during expectation of reward. A mice showed a reduced theta-gamma coupling in the NAc in comparison to ENA mice (unpaired t-test, *P<0.05). **P** PFC-NAc circuit communication (PSI) at lgamma during reward expectation.

After surgical recovery, electrophysiological recordings were first performed in standard cages immediately after the FR5 sessions. Locomotor activity was similar in both groups (Supplementary Fig.2A), and no significant differences in power were detected between addicted and extreme non-addicted mice, neither in the mPFC nor the NAc (Supplementary Fig.2B). Notably, both groups exhibited a flow of information (phase slope index, PSI) from the NAc to the mPFC at theta and hgamma frequencies (NAc to mPFC PSI, NAc leads) and a mPFC to NAc flow of information at lgamma frequencies (mPFC to NAc PSI, mPFC leads) under basal conditions (Supplementary Fig.2C). However, no significant differences were found in circuit communication between addicted and extreme non-addicted mice.

We next recorded neural activities during FR5 post-surgery for 10 FR5 sessions, where mice of both groups reduced the number of reinforcers per session from 51.16 to 41.18 (Fig.1A). Addicted mice obtained more reinforcers and performed more lever-presses, exhibiting increased impulsivity and persistence to obtain rewards in comparison with extreme non-addicted mice (Fig.2B-E and Supplementary Figs.3 and 4A-C). We investigated the neurophysiological fingerprints of decision-making and reward expectation during these FR5 sessions (Supplementary Fig.5A-B). We defined decision-making epochs as the second prior to the first of the five lever-presses (Fig.2G). During decision-making epochs, oscillatory power in the mPFC and the NAc was similar between extreme non-addicted and addicted mice (Fig.2H). Cross-frequency coupling was also similar between phenotypes (Fig.2I and Supplementary Fig.6A-D). In contrast, circuit communication at hgamma and hfo were different in the two subgroups of mice. Extreme non-addicted mice displayed NAc to mPFC signaling at hgamma frequencies and mPFC to NAC signaling at high frequencies. These electrophysiological signatures were significantly disrupted in addicted mice (Fig.2J; Supplementary Fig.6E). Notably, a negative correlation was detected between the number of reinforcers and the mPFC to NAc hfo circuit communication when including both phenotypes (Fig.2K), but not between NAc to mPFC hgamma communication. These findings suggested that mPFC-NAc connectivity was disrupted in addicted mice during decision-making, particularly at hfo ranges, whereas local neural synchrony was unaffected.

The differential neural substrates of reward expectation were also investigated in addicted and extreme non-addicted mice. We defined reward expectation epochs as the second occurring immediately after the fifth lever press when the cue light turned on and reward was delivered (Fig.2L). mPFC power was similar between phenotypes during reward expectation. However, addicted mice showed decreased power at lgamma frequencies in the NAc compared to extreme non-addicted mice (Fig.2M). Notably, lgamma power in the NAc correlated negatively with the number of reinforcers when including both genotypes (Fig.2N). Moreover, theta-gamma coupling was reduced in the NAc of addicted animals compared to extreme non-addicted mice (Fig.2O). In terms of circuit communication, we identified a mPFC to NAc signaling predominance at lgamma, hgamma and hfo (mPFC to NAc PSI, mPFC leads) (Fig.2P and Supplementary Fig.7F) that was similar in both groups. These results indicate that during reward expectation addicted mice displayed disrupted lgamma activity locally in the NAc, while mPFC synchrony and circuit connectivity remained unaltered (Supplementary Fig.7A-F).

### Addicted mice exhibit abnormal prefrontal-accumbens neural dynamics during active rewarding periods

We next investigated neural activity in the mPFC and NAc during the different rewarding and non-rewarding periods of the FR5 sessions. Changes between phases were analyzed in two seconds epochs (last second of the active period 1 to first second of the non-rewarding period and last second of the non-rewarding period to first second of the active period 1) (Fig.3A).

**Fig. 3.**
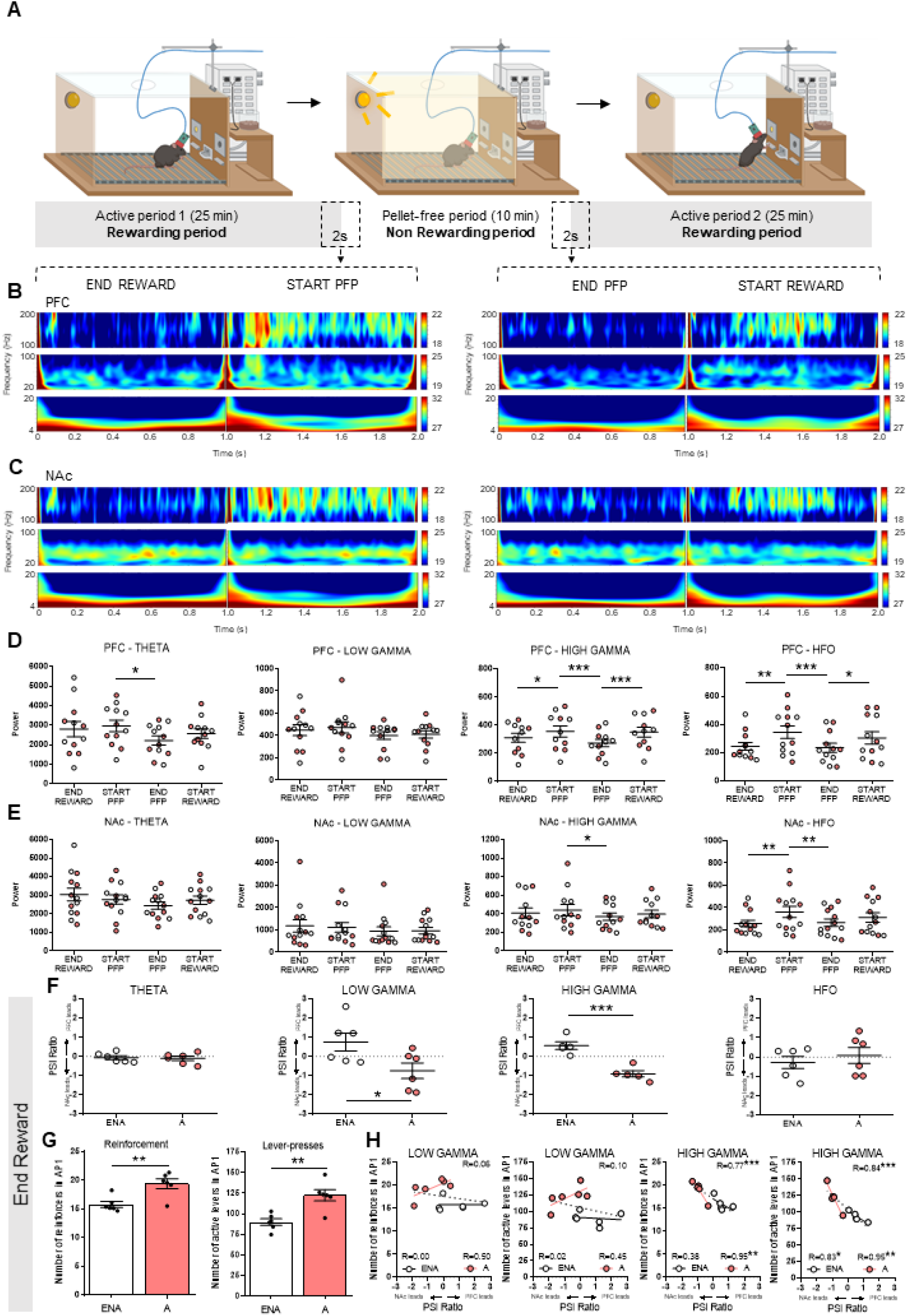
Addicted mice exhibit abnormal prefrontal-accumbens neural dynamics during active rewarding periods. **A** Schematic diagram illustrating the different phases of an FR5 operant session. The self-administration sessions are composed of two pellet periods (active periods) of 25 minutes separated by a pellet-free period of 10 min. During the pellet periods, pellets are delivered contingently after five active lever-presses paired with a cue light, whereas, during the pellet-free period, no pellet is delivered being this period signaled by the illumination of the entire self-administration chamber. Changes between phases were analyzed in two seconds epochs (1 second end reward to 1 second start pellet-free period and 1 second end pellet-free period to 1 second start reward). **B-C** Spectrograms of neural signals during the change between phases of the operant session (end reward to start PFP and end PFP to start reward) in the **B** mPFC and the **C** NAc. The plots have been divided into 4-20, 20-100 and 100-200 Hz to facilitate the comparison. **D-E** Power quantification during phase changes of the operant session at theta, lgamma, hgamma and hfo in the **D** mPFC and the **E** NAc. Extreme non-addicted (ENA) mice correspond to white colored dots whereas A mice are represented with red colored dots. Data are represented as mean ±SEM (one-way ANOVA repeated measures, *P<0.05, **P<0.01, ***P<0.001). **F** PFC-NAc circuit communication (PSI) at theta, lgamma, hgamma and hfo frequencies in ENA and A mice during the end of the rewarding period (end reward) (unpaired t-test, *P<0.05, ***P<0.001). **G** Behavioral data of reinforcement levels and number of lever-presses in the first active period (AP1) (unpaired t-test, **P<0.01). **H** Pearson correlations between mPFC to NAc circuit communication (PSI) at lgamma and hgamma bands; and behavioral data (number of reinforcers and lever-presses) acquired in the AP1 of the FR5 operant sessions. PFC-NAc communication at hgamma correlated with behavioral data in ENA and A mice together (marked with black dotted line) and separately (marked with a black line for ENA mice and with a red line for A mice) (*P<0.05, **P<0.01, ***P<0.001).

The start of the non-rewarding period was characterized by prominent increases in hgamma and hfo in the mPFC while hfo were also increased in the NAc (Fig.3B-E and Supplementary Fig.8A,B). Remarkably, at the end of the non-rewarding period delta, hgamma, and hfo power, decreased in the mPFC and NAc with respect to the beginning of this period (Fig.3D-E and Supplementary Fig.8A,B). Theta and beta oscillations were also reduced in the mPFC at the end of the non-rewarding period (Fig.3D and Supplementary Fig.8A). Subsequently, at the beginning of the second active period, mPFC power at hgamma and hfo increased in comparison to the end of the non-rewarding period (Fig.3D). These power fluctuations across different phases of the FR5 operant sessions indicated that neural activities at hgamma and hfo frequencies in the mPFC and NAc might contributed to encode relevant behaviors linked to persistence and reward, although similar changes were detected in addicted and extreme non-addicted mice (Fig.3D-E).

Furthermore, circuit communication analyses indicated a flow of information at lgamma and hgamma from mPFC to NAc in extreme non-addicted mice (mPFC to NAc PSI, mPFC leads) during active rewarding periods (end reward) that were reversed in addicted mice (NAc to mPFC PSI, NAc leads, Fig.3F). These observations suggest that sensitivity to reward might be encoded by lgamma and hgamma mPFC to NAc connectivity. Fluctuations within mPFC-NAc circuits were gamma specific and did not occur at other bands (Fig.3F and Supplementary Fig.8C). Correlation between circuit communication (PSI) and reward intake during the first rewarding period (active period 1), were also examined. As expected, during the first rewarding period addicted mice showed more rewards and lever-presses than extreme non-addicted mice (Fig.3G). We found that mPFC-NAc communication at hgamma, but not lgamma, correlated inversely with the number of reinforcers and active lever-presses when including both extreme genotypes (Fig.3H). These correlations were even stronger for the addicted group alone, where NAc to mPFC hgamma directionality was predominant. No differences in circuit communication were found in the other session periods analysed (Supplementary Fig.8D-F). Consistent with the literature^14,18^, these evidences suggest that the NAc of addicted mice governed over the NAc-mPFC circuit during rewarding periods.

### CB1 receptors shape prefrontal–accumbens neural dynamics

We investigated the acute effects of CB1 antagonism on behavior and prefrontal-accumbens neural dynamics of addicted and extreme non-addicted mice. First, the selective CB1 antagonist, rimonabant (3mg/kg) was acutely administered to mice freely moving in their home cages. Electrophysiological recordings were performed to study the time-course of rimonabant’s effects on prefrontal-accumbens neural dynamics. 1, 16, and 19 minutes after its administration in addicted and extreme non-addicted mice. Rimonabant induced modest alterations of neural oscillations within the mPFC and NAc and did not affect locomotor activity (Supplementary Fig.9A). Differences in power were observed between control and treated animals (addicted and extreme non-addicted combined) in the mPFC and the NAc 19 minutes after rimonabant administration (Fig.4A-C). Rimonabant progressively increased the delta band in both the mPFC and the NAc, and lgamma band in the NAc (Fig.4B-D) of both phenotypes compared to the vehicle. The rest of the frequency bands followed a similar power pattern compared to the vehicle in both areas (Supplementary Fig.9B).

**Fig. 4.**
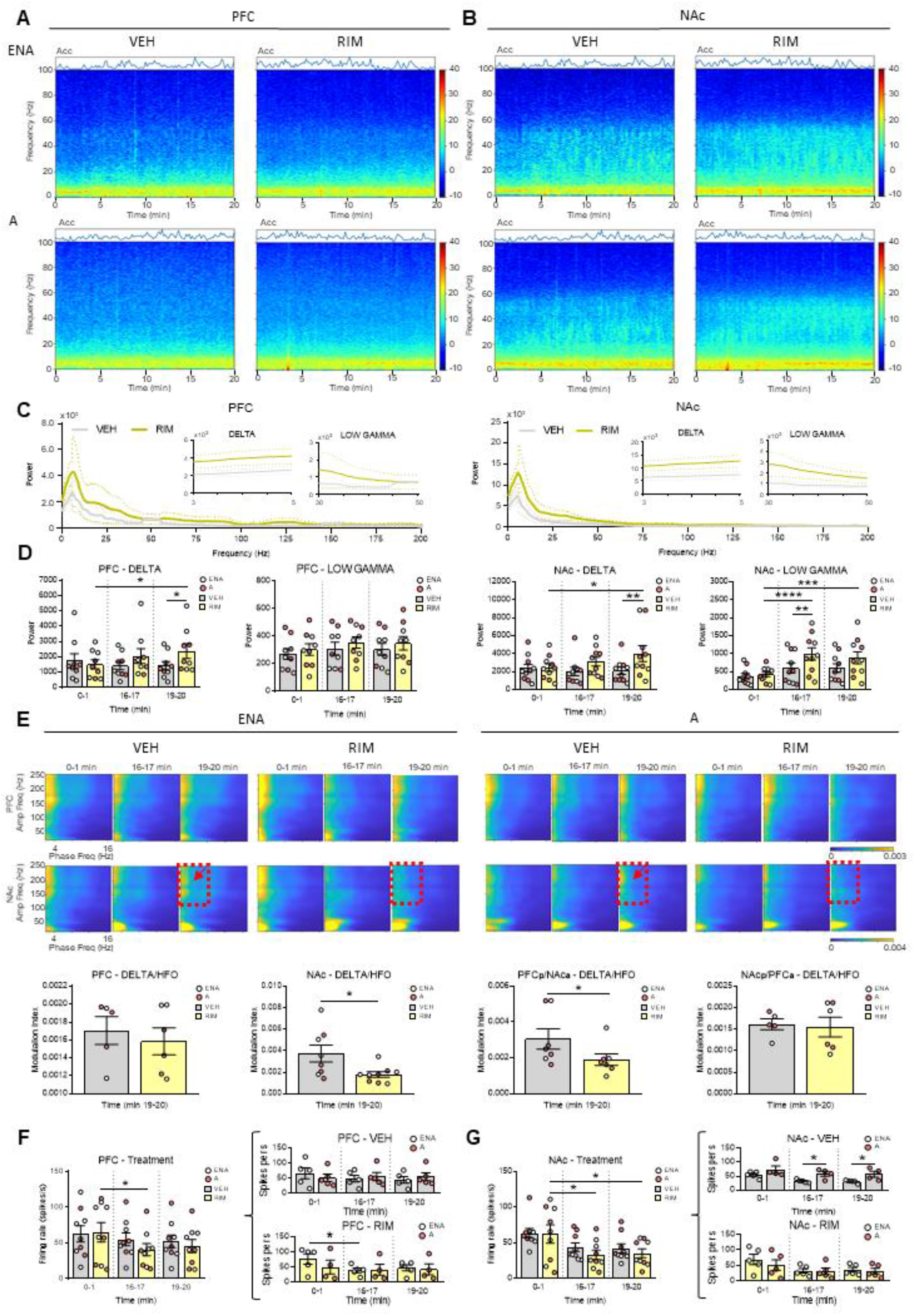
CB1 receptors shape prefrontal–accumbens neural dynamics. **A-B** Averaged spectrograms of signals of extreme non-addicted (ENA) and A mice under vehicle and rimonabant administration in the **A** mPFC and the **B** NAc. Panels represent the raw data with the corresponding quantification of the animals motility (accelerometer, variance of signals from the accelerometer integrated within headstages). **C** Power spectra of mPFC and NAc signals at the 19^th^ minute after administration of vehicle (grey) or rimonabant (yellow). Power values amplification of delta and lgamma are also shown. **D** Corresponding power quantification of delta and lgamma frequencies across time (0-1, 16-17 and 19-20 minutes after administration of vehicle or rimonabant) in the mPFC and the NAc. ENA mice are represented with white colored dots whereas A mice are represented with red colored dots. Differences are reported as mean ±SEM, two-way ANOVA multiple comparisons, *P<0.05, **P<0.01, ***P<0.001, ****P<0.0001. **E** Comodulation maps and quantification of local cross-frequency coupling in the mPFC and the NAc in 1 minute periods (0-1, 16-17 and 19-20 min) in ENA and A mice. The x-axis represents phase frequencies (2-16 Hz) and the y-axis represents amplitude frequencies (10–250 Hz). Numbers on top indicate the minute after vehicle or rimonabant administration. The red dotted square indicates the coupling between delta and hfo present in normal conditions (vehicle), marked with a red arrow, that disappear after 19 minutes of rimonabant administration (paired t-test, *P<0.05). **F-G** Time-course of changes in the firing rate of neurons under vehicle and rimonabant treatment in the **F** mPFC and the **G** NAc. Multi-unit activity was used in this analysis. Differences are reported as mean ±SEM, two-way ANOVA multiple comparisons, *P<0.05.

We further assessed circuit communication and cross-frequency coupling under rimonabant. Circuit communication, measured with PSI was unaffected by rimonabant (Supplementary Fig.9C). In the mPFC, delta-hfo coupling was detected, although no significant differences were observed among phenotypes or treatments. In the NAc, addicted and extreme non-addicted mice exhibited robust delta-lgamma and delta-hfo coupling with vehicle. The delta-hfo (1-4Hz with 150-250Hz) coordination subsided 19 minutes after rimonabant administration (Fig.4E). At circuit level, rimonabant effects were similar to those found in the NAc. The phase of mPFC delta modulated the amplitude of NAc hfo (PFC_phase_-NAc_amplitude_).

As observed in the NAc, rimonabant weakened this circuit coupling (Fig.4E and Supplementary Fig.9D). By contrast, no significant changes were detected for the reversed inter-regional coupling, NAc_phase_-PFC_amplitude_ between vehicle and rimonabant treated groups (Fig4E and Supplementary Fig.9D). These findings demonstrate that prefrontal-accumbens communication is sensitive to CB1 pharmacological blockade under basal conditions.

We also examined the effects of rimonabant on the firing rate of neuron populations under basal conditions. Rimonabant decreased population activity in the mPFC and NAc considering both groups of mice. Furthermore, firing rates in the mPFC slightly decreased 16 minutes after rimonabant in extreme non-addicted mice, but not in addicted mice (Fig.4F). After vehicle, treat-addicted mice showed higher neuron population activity in the NAc than extreme non-addicted mice, an effect rescued by rimonabant (Fig.4G). Therefore, power, cross-frequency coupling and spiking activity in prefrontal-accumbens circuits of both phenotypes were sensitive to CB1 receptor blockade.

### Rimonabant reduces reinforcement levels and reverses abnormal prefrontal-accumbens neural dynamics during decision-making and reward expectation

We used rimonabant to investigate the neural correlation underlying its therapeutic effects on food addiction. The treatments were alternated for four consecutive days (Fig.1A). As reported in previous studies^35,36^, CB1 antagonism strongly reduced reinforcement responding. The number of reinforcers drastically decreased in both phenotypes in comparison with the vehicle groups (Fig.5A left and Fig.5A right): 25.5% of basal levels in addicted mice and close to 0 in extreme non-addicted mice (Fig.5B). The number of reinforcers consumed in addicted mice when treated with rimonabant correlated inversely with NAc lgamma power recorded immediately prior to the operant session (Fig.5C). No correlation was detected between the decreased reinforcement levels and delta activity, neither in the mPFC nor in the NAc (Supplementary Fig.10A). The fact that extreme non-addicted animals consumed a minimum number of reinforcers under rimonabant treatment limited the electrophysiological analysis. Therefore, we focused our analyses on the electrophysiological effects of rimonabant in addicted mice.

**Fig. 5.**
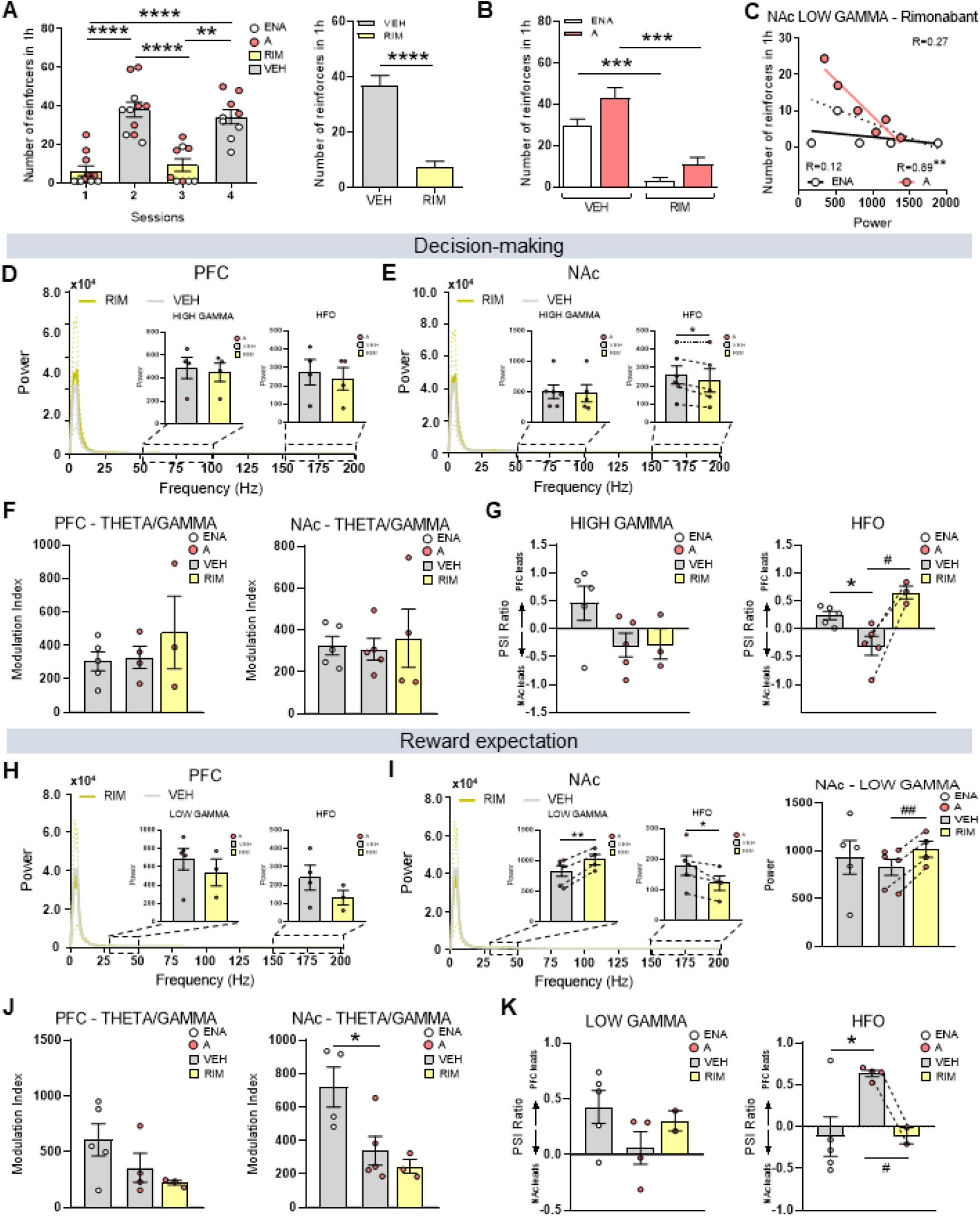
Rimonabant reduces reinforcement levels and reverses abnormal prefrontal-accumbens neural dynamics during decision-making and reward expectation. **A** Reinforcement values of extreme non-addicted (ENA) and A mice under vehicle and rimonabant treatment. ENA mice are represented with white dots whereas A mice are shown with red dots. Grey bars indicate vehicle treatment and yellow bars indicate rimonabant treatment. In the left, number of reinforcers in 1h operant session during 4 consecutive days alterning vehicle and rimonabant administration. In the right, corresponding mean of reinforcement levels of the 2 sessions under vehicle and rimonabant administration assembling ENA and A mice together. Differences are reported as mean ±SEM, One-way ANOVA repeated measures, **P<0.01, ****P<0.0001 and Paired t-test, ****P<0.0001. **B** Corresponding mean of reinforcement levels in ENA and A mice after vehicle and rimonabant administration. Data are represented as mean ±SEM, Paired t-test, ***P<0.001. **C** Pearson correlation between the number of reinforcers in rimonabant treatment FR5 sessions and NAc lgamma power increase observed after 19 minutes of rimonabant administration (**P<0.01). The dotted line corresponds with linear regression of ENA and A mice together, whereas linear regressions in ENA and A mice separately are represented in black and red colored lines, respectively. **D-E** Power spectra of **D** mPFC and **E** NAc signals during decision-making after administration of vehicle (grey) or rimonabant (yellow). Power values amplification of hgamma and hfo are also shown (paired t-test, *P<0.05). **F** Quantification of local theta-gamma coupling in the mPFC and NAc during decision-making (1s) in ENA and A mice treated with vehicle and rimonabant treated A mice. **G** PFC-NAc circuit communication (PSI) at hgamma and hfo frequencies during decision-making under vehicle and rimonabant treatment. Data are represented as mean ±SEM, unpaired and paired t-test, *P<0.05, #P<0.05, respectively. **H-I** Power spectra of **H** mPFC and **I** NAc signals during expectation of reward after administration of vehicle (grey) or rimonabant (yellow). Power values amplification of lgamma and hfo are also shown. The corresponding power quantification of lgamma in the NAc is also represented in the right with the ENA group treated with vehicle (paired t-test, ##P<0.01). **J** Quantification of local theta-gamma coupling in the mPFC and NAc during reward expectation (1s) in ENA and A mice treated with vehicle and rimonabant treated A mice (unpaired t-test, *P<0.05). **K** PFC-NAc circuit communication (PSI) at lgamma and hfo frequencies during reward expectation under vehicle and rimonabant treatment (unpaired and paired t-test, *P<0.05, #P<0.05).

During decision-making periods, mPFC power of addicted mice did not vary when treated with rimonabant (Fig.5D) while the power of hfo slightly decreased in the NAc (Fig.5E). Moreover, theta-gamma coupling in both areas was similar in both phenotypes and not overtly affected by the treatment (Fig.5F and Supplementary Fig.10B). We previously identified that NAc-to-mPFC hgamma and mPFC-to-NAc hfo signals were involved in decision-making in extreme non-addicted mice and were altered in addicted mice (Fig.2J-K). At high frequencies, the addicted vehicle control group displayed again similar reversed directionality of signals compared with extreme non-addicted mice (NAc to mPFC PSI, NAc leads) (Fig.5G). In rimonabant-treated addicted mice the directionality of signals reversed resembling those of extreme non-addicted animals (mPFC to NAc PSI, mPFC leads) (Fig.5G). These power and connectivity results revealed that rimonabant corrected aberrant signals at high frequencies during decision-making periods in addicted animals. Neural activities at other frequencies were unchanged (Supplementary Fig.10B).

We also studied the electrophysiological fingerprints of CB1 antagonism during reward expectation periods. No significant changes of mPFC power were detected in rimonabant-treated addicted mice in comparison with the vehicle group (Fig.5H). In the NAc, lgamma power was larger in rimonabant-treated addicted mice than in vehicle groups (Fig.5I). Furthermore, as during decision-making epochs, we detected a power reduction at hfo in rimonabant-treated mice (Fig.5I). Theta-gamma coupling was again reduced in the NAc of addicted animals during expectation of reward (Fig.2O), although it was not rescued by rimonabant (Fig.5J). Moreover, the mPFC-to-NAc lgamma signals and NAc-to-mPFC hfo signals observed in extreme non-addicted were disrupted in addicted animals. Thus, hfo traveled in the opposite direction (from the mPFC to the NAc) and these signals were rescued by rimonabant (Fig.5K). The remaining frequency bands were unaltered by rimonabant during reward expectation periods (Supplementary Fig.10E).

## Discussion

This study describes for the first time neurophysiological signatures within prelimbic mPFC– NAc core circuits underlying behavioral responses associated with food addiction that are rescued by the blockade of CB1, cannabinoid receptors. Using a behavioral paradigm with high translational face validity to human addiction^27,36^, we mimicked the development of addiction after repeated seeking of palatable food in an operant training during 110 sessions. This mouse model allowed to classify mice into vulnerable (29.03%) or resilient (70.97%) to develop food addiction. We further identified a subgroup of resilient mice with extreme non-addicted phenotypes (20%). These percentages are consistent with those obtained in previous studies using the same mouse model^27,36^.

Homecage recordings performed following the FR5 operant conditioning sessions revealed that local synchrony (power and cross-frequency coupling) within the mPFC and NAc and the circuit communication were similar between extreme non-addicted and addicted mice under basal conditions. Under these conditions, we identified flows of information from the NAc to the mPFC at theta and hgamma frequencies and a mPFC to NAc flow at lgamma frequencies in both phenotypes that unraveled a rich communication between both regions.

Consistently, theta and gamma rhythms have been reported to participate in mechanisms underlying rewarding responses^37,38^. The predominance of the NAc to mPFC theta and hgamma signaling observed here might represent the prolonged activation of the NAc by the DA system triggered by the consumption of palatable reinforcers during the food-seeking sessions^9^. Moreover, lgamma signals traveling from the mPFC to the NAc might correspond to mPFC-dependent executive functions required for the proper performance of the task^39^.

To determine the neural correlates underlying the distinct phenotypes observed, we focused on two main behaviors contributing to reward seeking, the main variable identified by PCA to contribute to food addiction behaviors: decision-making and reward expectation. Both, altered decision-making and outweighed reward expectation have been directly linked with reinforcing processes of food consumption^40,41^. Our findings revealed that mPFC–NAc circuit communication at high frequencies (hgamma and hfo) was disrupted in addicted mice during decision-making, whereas local synchrony was unaltered. The directionality of hgamma signaling in addicted mice (prelimbic mPFC to NAc core) was the opposite to that observed in extreme non-addicted mice (NAc core to prelimbic mPFC). Notably, mPFC to NAc hfo signals detected in extreme non-addicted mice during decision-making were disrupted in addicted animals (PSI close to cero). The fact that this neural marker correlated with the number of reinforcers support the notion of a top-down dysregulation of executive function that favored addictive-like behaviors. Gamma and hfo have been associated with attention, being these frequency bands recognized as potential biomarkers in cognitive disorders^42,43^.

Additionally, it is well known that cognitive impairment led by alterations in the mPFC and the hippocampus are closely associated with poor decision-making^44–46^, which is also a characteristic of addiction^47,48^. Thus, these previous results support our findings suggesting that alterations of hgamma and hfo during decision-making may participate in the expression of behavioral anomalies linked to food addiction. Coordination at hfo may be characteristic of NAc microcircuits^49^. These rhythms are generated in the NAc and are used by this area to exchange information with other brain areas, including the mPFC^50^. In support of this, previous studies from our group and others have not detected communication in mPFC-hippocampus circuits at those high frequencies^51–53^. Thus, our results highlight the role of synchrony at high frequencies in the mesocorticolimbic pathway during reward-seeking behaviors that may be disturbed in food addiction.

During reward expectation, we found that addicted mice showed local desynchronization of gamma rhythms in the NAc that correlated with reinforcement levels. Moreover, addicted mice presented weakened theta-gamma coupling in the NAc. Together, these findings put forward a relevant role for local gamma synchrony within NAc microcircuits in the pathological reward expectation occurring during food addiction. The NAc plays crucial roles in reward prediction during reward-seeking behavior^54^, and changes in theta and gamma oscillations have been reported in the NAc and mPFC areas^55,56^. DA signals in the NAc are crucial for obtaining and consuming drug or food rewards^14^ and addiction is associated with a decrease in DA release in response to drugs of abuse^17^. Therefore, we suggest that disruption of lgamma and theta-gamma coupling in addicted mice may result from decreased DA release in the NAc, and the consequent alterations induced in the encoding of reward prediction.

The contribution of gamma and hfo rhythms within PFC-NAc circuits to reward-seeking was further supported by fluctuations of power during the different phase changes of the operant sessions. Thus, oscillatory activities >50Hz (hgamma and hfo) increased at the start of the non-rewarding period, pointing to their involvement in persistence of response since mice are still responding during this period in spite of the absence of reward. This was further supported by the reduction of these oscillatory activities at the end of the non-rewarding period, presumably when persistence of response had declined.

Furthermore, addicted mice exhibited NAc to mPFC lgamma and hgamma signaling at the end of the first rewarding period, opposite to the mPFC to NAc flow of information observed in extreme non-addicted mice. Remarkably, abnormal hgamma circuit communication correlated strongly with increased food intake and lever-presses in addicted mice. Previous studies have established that the drug-induced alterations of DA transmission impairs top-down processes controlled by the mPFC^18^, suggesting that the inability to stop consuming drugs suffered by patients with addiction may be due to a failure of inhibitory control^19,57^. Therefore, we hypothesize that this altered flow of information reflects the disruption of mPFC function leading to the failure of inhibitory control in addicted mice, which promotes persistence in seeking rewards.

Reinforcing properties of palatable foods depend on DA release in the mesolimbic pathway that is modulated by the endocannabinoid system^58,59^. Thus, CB1 receptor activation promotes DA release in the NAc by decreasing GABAergic inhibitory synaptic inputs to DA neurons^34^. In basal conditions, rimonabant augmented delta (~5Hz) oscillations in the mPFC and NAc and lgamma oscillations in the NAc, while not modifying locomotor mobility in any groups of animals. Concomitantly, NAc and inter-regional delta-hfo coordination weakened after rimonabant administration. Interestingly, addicted animals treated with vehicle exhibited modest increases in NAc spiking activity compared to extreme non-addicted mice and this activity was rescued by rimonabant administration. Rimonabant also decreased spiking activity in the mPFC of all the mice, but did not affect the directionality of signals within the circuit. Overall, the effects of rimonabant on the intrinsic neural dynamics of mPFC-NAc circuits during basal conditions were modest pointing to an involvement of CB1 receptors in the modulation of neural activities encoding specific behaviors, but not in the homeosthasis of the circuit. In agreement, rimonabant exerted strong effects on behavioral responses and neural activities associated with food addiction during the operant sessions. Indeed, it dramatically reduced the number of reinforcers obtained by addicted and extreme non-addicted mice. This decrease correlated inversely with NAc lgamma power recorded previously in the homecage. This finding implicated once again disrupted gamma synchrony in the NAc as a possible mechanism underlying pathological food seeking. Notably, rimonabant increased lgamma oscillations in the NAc prior to the operant conditioning in all mice, suggesting a role of CB1 antagonism in the neural substrates underlying the amelioration of addictive-like behaviors.

The effects of rimonabant on neural activities associated with decision-making and reward expectation during food-seeking behaviors also provided strong support for this hypothesis. First, during decision-making, rimonabant restored mPFC control of the circuit in addicted mice. In fact, the directionality of signals reversed consistently in all rimonabant-treated addicted mice resembling those of extreme non-addicted animals. Second, during reward expectation periods, both lgamma power in the NAc and the circuit’s directionality at high frequencies were normalized in addicted mice to extreme non-addicted levels. Previous studies have proposed that CB1 receptors contribute to the dynamic coordination of prefrontal-hippocampal circuits^60^. Here, we extend these observations and demonstrate a modulation of mPFC-NAc pathways by CB1 receptors in mice during behaviors linked to food addiction.

Altogether, we identified specific neural signatures of PFC-NAc circuits in food-addicted mice during active rewarding periods as well as during decision-making and reward expectation. We further demonstrate the rescue of most of these abnormal behaviors and neural activities by the blockade of CB1 receptors. Our results may elucidate the neural substrates underlying food addiction and eating disorders, shedding light into the understanding of these pathologies and the identification of novel therapeutic targets.

## Acknowledgements

We thank J. De la Rocha, J. Santos, M. Alemany-González, M. Linares, R. Martín, D. Real, F. Porrón and F. Geraci for helpful comments and technical support to this study. We also thank Nicholas A. Donnely for providing the matlab script to analyze 1second PAC. This work was supported by the Spanish “Minsterio de Ciencia e Innovación (MICIN), Agencia Estatal de Investigación (AEI)” (PID2020-120029GB-I00/MICIN/AEI/10.13039/501100011033, RD21/0009/0019, to RM; the “Generalitat de Catalu-nya, AGAUR” (2017 SGR-669, to RM; the “ICREA-Acadèmia” (2020, to RM); the “European Commission-DG Research” (PainFact, H2020-SC1-2019-2-RTD-848099, QSPain Relief, H2020-SC1-2019-2-RTD-848068, to RM); the Spanish “Instituto de Salud Carlos III, RETICS-RTA” (RD16/0017/0020, to RM); the Spanish “Ministerio de Sanidad, Servicios Sociales e Igualdad, Plan Nacional Sobre Drogas” (PNSD-2021I076, to RM; PNSD-2019I006, to EMG; Ministerio de Ciencia e Innovación (ERA-NET) PCI2021-122073-2A to EMG. This work was supported by the Spanish Research Agency (Agencia Estatal de Investigación, AEI/FEDER) project PID2019-104683RB-I00 to M.V.P.

## Author Contribution

E.M-G., M.V-P., and R.M. conceived and designed the behavioral and electrophysiological studies; P.C. performed the behavioral experiments and the statistical analyses and graphs with the collaboration of L.R. and the supervision of E.M-G., M.V-P., and R.M.; T.G. performed the surgeries. S.K. collaborated with the behavioral experiments. P.N. and P.C. performed the bioinformatic analysis supervised by M.V.P; P.C., E.M-G., M.V.P. and R.M. wrote the manuscript with inputs from all the other authors.

## Additional information

**Supplementary information** accompanies this paper.

**Correspondence** and requests for materials should be addressed to R.M.

