## Supplementary material for "Prefrontal–accumbens neural dynamics abnormalities in mice vulnerable to develop food addiction": SI

Calvé, P et al.

**Methods**

**Animals.** Experiments were performed in C57BL/6J male mice (n=31). Mice (2 - 10 months old) were housed under conditions of controlled temperature (23 ±1ºC) and illumination (12 hours light/dark cycle). All procedures were conducted in compliance with EU directive 2010/63/EU and Spanish guidelines (Laws 32/2007, 6/2013 and Real Decreto 53/2013) and were authorized by the local ethical committee (Comitè Ètic d'Experimentació Animal-Parc de Recerca Biomèdica de Barcelona, CEEA-PRBB, agreement N°10554).

**Behavioral experiment.** Daily self-administration sessions maintained by chocolate-flavored pellets were performed in operant boxes (Med Associates Inc) and lasted 1 hour. The beginning of each self-administration session was signaled by turning on a house light placed on the ceiling of the skinner box during the first 3 seconds. The self-administration sessions were composed of two pellet periods (active periods) of 25 minutes separated by a pellet-free period of 10 min. During the pellet periods, pellets were delivered contingently after an active response paired with a stimulus light (cue light). A time-out period of 15 seconds was established after each pellet delivery, where the cue light was off, and no reinforcer was provided after responding on the active lever. Responses on the active lever and all the responses performed during the time-out period were recorded. During the pellet-free period, no pellet was delivered and this period was signaled by the illumination of the entire self-administration chamber. In the operant conditioning sessions, mice were under fixed ratio 1 (FR1) schedule of reinforcement (one lever-press resulted in one pellet delivery) followed by an increased FR to 5 (FR5) (five lever-presses resulted in one pellet delivery) for the rest of the sessions. The criteria for the achievement of the operant responding were acquired when all of the following conditions were met: (1) mice maintained a stable responding with <20% deviation from the mean of the total number of reinforcers earned in three consecutive sessions (80% of stability); (2) at least 75% responding on the active lever; and (3) a minimum of 5 reinforcers per session. After each session mice were returned to their home cages. Three behavioral tests were used to evaluate the food addiction-like criteria as recently described and adapted from cocaine addiction-like in rats (Deroche-Gamonet et al., 2004; Deroche-Gamonet and Piazza, 2014). These three criteria summarized the major hallmarks of addiction based on DSM-IV, specified in DSM-5 and now included in the food addiction diagnosis through the YFAS 2.03. (1) Persistence of response: Non-reinforced active responses during the pellet-free period (10 min), when the box was illuminated and signaling the unavailability of pellet delivery, were measured as persistence of food-seeking behavior. On the three consecutive days before the progressive ratio, mice were scored. (2) Motivation: The progressive ratio schedule of reinforcement was used to evaluate the motivation for the chocolate-flavored pellets. The response required to earn one single pellet escalated according to the following series: 1, 5, 12, 21, 33, 51, 75, 90, 120, 155, 180, 225, 260, 300, 350, 410, 465, 540, 630, 730, 850, 1000, 1200, 1500, 1800, 2100, 2400, 2700, 3000, 3400, 3800, 4200, 4600, 5000, and 5500. The maximal number of responses that the animal performs to obtain one pellet was the last event completed, referred to as the breaking point. The maximum duration of the progressive ratio session was 5 h or until mice did not respond on any lever within 1 h. (3) Compulsivity: Total number of shocks in the session of shock test (50 min) performed after the PR test, when each pellet delivered was associated with a punishment, were used to evaluate compulsivity-like behavior. Mice were placed in a self-administration chamber without the metal sheet with holes and consequently with the grid floor exposed (contextual cue). In this shock-session, mice were under a FR5 schedule of reinforcement during 50 minutes with two scheduled changes: at the fourth active lever-response mice received only an electric footshock (0.18 mA, 2 seconds) without pellet delivery and at the fifth active lever-response, mice received another electric footshock with a chocolate-flavored pellet paired with the cue light. The schedule was reinitiated after 15 seconds pellet delivery (time-out period) and after the fourth response if mice did not perform the fifth response within a min. After performing the three behavioral tests to measure the food addiction-like behavior, mice were categorized in two extreme subpopulations (addicted or extreme non-addicted animals) depending on the number of positive criteria that they had achieved. An animal was considered positive for an addiction-like criterion when the score of the specific behavioral test was above the 75th percentile of the normal distribution. Mice that achieved two or three addiction-like criteria were considered as addicted animals and mice that achieved zero or one addiction-like criterion were considered as extreme non-addicted animals.

**Surgeries and electrode implantation**. Mice (n=18) were induced with a mixture of ketamine/xylazine and placed into a stereotaxic apparatus for intracranial electrode implantation. Anesthesia was maintained with isoflurane 0.25 - 2%. Micro-screws were placed into the skull to stabilize the implant. An additional micro-crew was placed on the top of the cerebellum and used as a general ground. Two tungsten electrodes (25 um wide; 100 to 400 kOhm; Advent, UK) were unilaterally implanted in the prelimbic mPFC (AP: 1.98 mm, ML: 0.3 mm, DV: -2.30 mm) and in the NAc core (AP: 1.10 mm, ML: 1.0 mm, DV: -4.60 mm). Neural activity was recorded while the electrodes were being lowered down inside the brain. After surgery, animals were allowed one week to recover and habituate to the implant connected to the recording cable. After the end of the experiments, electrode placements were confirmed histologically by staining brain slices (30-μm-thick coronal slices using a cryostat) using Cresyl violet. Images were obtained and analyzed using light microscopy (Olympus BX61) with a 4x magnification. Electrodes with tips outside the targeted areas were discarded from data analyses.

**In vivo electrophysiological recordings.** Electrophysiological recordings were conducted in the dark illumination cycle (from 8:00 a.m. to 7:30 p.m.). Animals were removed from their home cages, connected to the headstage and placed inside the skinner box in order to start the session. All the recordings, local field potentials (LFPs) and MUA were carried out with the multi-channel Open Ephys system at 0.1-6000 Hz and a sampling rate of 30 kHz with Intan RHD2132 amplifiers equipped with an accelerometer. Two animals were recorded simultaneously in separated skinner boxes. To obtain LFPs, electrophysiological recordings were sampled to 1 kHz, decoded, and filtered with customized Python scripts. The frequency bands considered for specific band analyses were delta (1-4 Hz), theta (8-12 Hz), beta (15-25 Hz), low gamma (30-50 Hz), high gamma (50-100 Hz) and high frequency oscillations (150-200 Hz). MUA was estimated by first subtracting the raw signal from each electrode with the signal from a nearby referencing electrode to remove artifacts related to the animal's movement. Then, continuous signals were filtered between 450-6000 Hz with Python and thresholded at -3 sigma standard deviations with Offline Sorter v4 (Plexon Inc.). The criteria for the achievement of the operant responding during electrophysiological recordings were acquired when the total number of reinforcers earned in three consecutive sessions (more than 60% of stability) and a minimum of 5 reinforcers per session.

**Experimental design.** C57BL/6J mice (n=31) were trained in operant boxes for 110 days to develop food addictive-like behaviors[27,36]. The long-term operant conditioning started with a fixed ratio of one lever press to one pellet reward (FR1) schedule of reinforcement during five sessions and was followed by 105 sessions under FR5 (Fig. 1A). One mouse that did not achieve the acquisition criteria after day 33 was excluded from the study. Persistence of response, motivation and compulsivity were evaluated in the early (session 18) and late periods (session 107). Persistence of response was evaluated by the number of non-reinforced active responses during the pellet-free period. Motivation was defined by the breaking point obtained during the progressive ratio (PR) schedule. Compulsivity was evaluated by the number of active responses associated with an electric footshock delivery. After the behavioral evaluation during the late training period, mice were classified into two different phenotypes: addicted (covering two or three criteria) and non-addicted (covering zero or one criterion) mice. We ordered each mouse on a quantitative gradual addiction scale (Fig. 2A). For this scale, we considered the individual score values of the three addiction criteria and the two phenotypic traits as factors of vulnerability to addiction of each mouse in the two extreme subpopulations. A score above the 75th percentile punctuated 3 points, mice between 50-75th 2 points, percentiles 25-50th 1 point, and above the 25th percentile with 0 points. Next, non-addicted mice (0 criteria) with a score above the 75th percentile punctuated with 0 points, mice between percentiles 50-75th 1 point, between 25-50th with 2 points, and beyond the 25th percentile with 3 points. The obtained mouse score was multiplied by 5 in compulsivity, 4.5 in the persistence of response, 4 in motivation, 3 in the persistence of the last ten days, 2.5 in impulsivity, 2 in the impulsivity of the last ten days, 1.5 in reinforcers obtained during three days and 1 in reinforcers obtained last ten days. These extreme subpopulations (extreme non-addicted and addicted mice) underwent stereotaxic surgery to implant stereotrodes (two twisted electrodes) in the mPFC and the NAc. Following recovery from surgery, recordings were performed in operant boxes adapted for electrophysiology for short-term operant training (post-surgery FR5 sessions 1 to 10). Following these ten sessions with electrophysiological recordings, mice were treated with rimonabant 3 mg/kg to investigate the effects on the neural substrates of the addictive phenotype. Recordings in standard cages and recordings during four FR5 sessions (post-surgery FR5 sessions 11 to 14) were performed to study the electrophysiological effects of rimonabant in the mPFC and the NAc (Fig. 1A-B, Supplementary Fig. 1A).

**Data and statistical analysis.** All analyses were carried out with custom scripts programmed in Python. Recorded signals from each electrode were detrended, notch-filtered to remove power line artifacts (50 Hz) and decimated to 1kHz offline to obtain LFPs. Noisy electrodes were discarded by visual inspection from individual channel spectrograms. Power spectral density results were calculated using the multi-taper method from the spectral_connectivity package in Python (time-half-bandwidth product = 5, 9 tapers, 60s sliding time window without overlap; https://github.com/Eden-Kramer-Lab/spectral_connectivity). Spectrograms of cage recordings were constructed using consecutive Fourier transforms (scipy.signal.spectrogram function, 60s time window, no overlap, no detrend). A 1/f normalization was applied to power spectral density results, and power spectrograms were scaled to decibels for visualization purposes. The frequency bands considered for the band-specific analyses included: delta (2-5 Hz), theta (8-12 Hz), beta (18-25 Hz), low gamma (30-50 Hz), high gamma (50-100 Hz) and high-frequency oscillations (150-200 Hz). Spectrograms of change phases during FR5 operant sessions were constructed using stockwell specs, 1second time window. Phase-amplitude coupling (PAC) was measured with a Python implementation of the method described in Tort et al. (phase frequencies = [0, 15] with 1 Hz step and 4 Hz bandwidth, amplitude frequencies = [10, 250] with 5 Hz step and 10 Hz bandwidth; (Tort et al., 2008). The length of the sliding window was 300 seconds for the overview plots and 60 seconds for the quantifications, without overlap. PAC quantification results were obtained by averaging the values of selected areas of interest in the comodulograms. Moreover, we used the Canolty method implemented on (Onslow et al., 2011) for the PAC measurements of decision-making and reward expectation epochs periods (1second windows). Prefrontal-accumbens circuit communication was estimated via the phase slope index (PSI). In addition, we calculated the flow of information between areas with the phase slope index (PSI) with a Python translation of MATLAB's data2psi.m (epleng = 60 sec, segleng = 1 sec) from (Nolte et al., 2008).

**Reporting summary.** Further information on research design is available in the Nature Research Reporting summary linked to this article.

**Data availability**

Individual data points are graphed in the main and Supplementary Figures. In addition, all relevant data that support this study are available from the corresponding author to any interested researcher upon reasonable request.

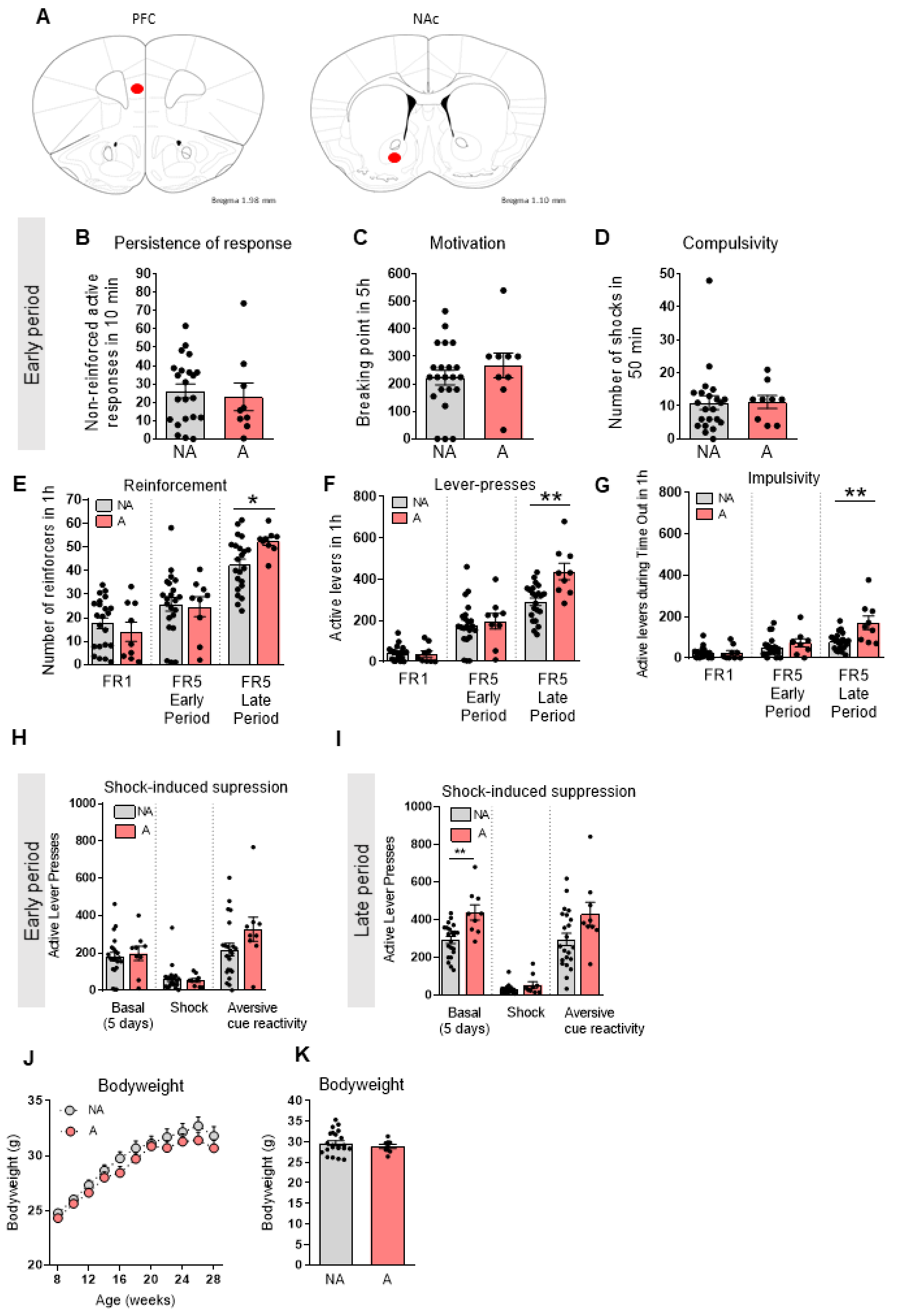


**Supplementary Fig. 1 Identification of extreme phenotypes of addicted and non-addicted mice by long-term operant training. A** Schematic diagram of the mPFC and NAc areas selected for electrode implantation. Two tungsten electrodes were unilaterally implanted in the prelimbic mPFC (AP: 1.98 mm, ML: 0.3 mm, DV: -2.30 mm) and in the NAc core (AP: 1.10 mm, ML: 1.0 mm, DV: -4.60 mm). (**B-D**) Behavioral tests of the three addiction-like criteria represented with the mean in both NA and A phenotypes at the early period. **B** Persistence to response. Total number of non-reinforced active responses during three consecutive daily 10-min of pellet-free period. **C** Motivation. Breaking point achieved in 5h of PR schedule. The breaking point refers to the maximal effort that an animal is willing to do to achieve one reward. **D** Compulsivity. Number of shocks that mice receive in 50 min in the shock test in which each pellet delivery was associated with a footshock (0.18 mA). **E-G** Behavioral data of phenotypic traits in FR5 sessions represented by individual values with the mean and ±SEM. FR1 corresponds to the five operant sessions of FR1, FR5 Early Period corresponds to the five FR5 sessions before the early period, FR5 Late Period corresponds to the five FR5 sessions before the late period. **E** Reinforcement levels quantified by counting the number of reinforcers consumed in 1h operant sessions. **F** Number of total active lever-presses scored in 1h operant session. **G** Impulsivity. Number of non-reinforced active lever-presses during five consecutive daily time out (15 s) after each pellet delivery (U Mann-Whitney, *P < 0.05, **P < 0.01). **H-I** Behavioral test of shock-induced suppression at the **H** early period and **I** late period represented by individual values with the mean of active levers and ±SEM. Basal corresponds to the five FR5 sessions before the compulsivity test. Shock corresponds to the compulsivity test where each pellet delivery was associated with a punishment, an electric foot shock. Shock-associated cue corresponds to the number of non-reinforced active responses in 50 min in the following session after the shock test with the same discriminative stimulus (grid floor) as shock test in which pressing the active lever had no consequences: no shock, no reward delivery and no cue light (U

Mann-Whitney, **P < 0.01). (J-K) Bodyweight. **J** Measurements of body weight in grams across weeks in NA and A mice. **K** Averaged body weight data per phenotype.

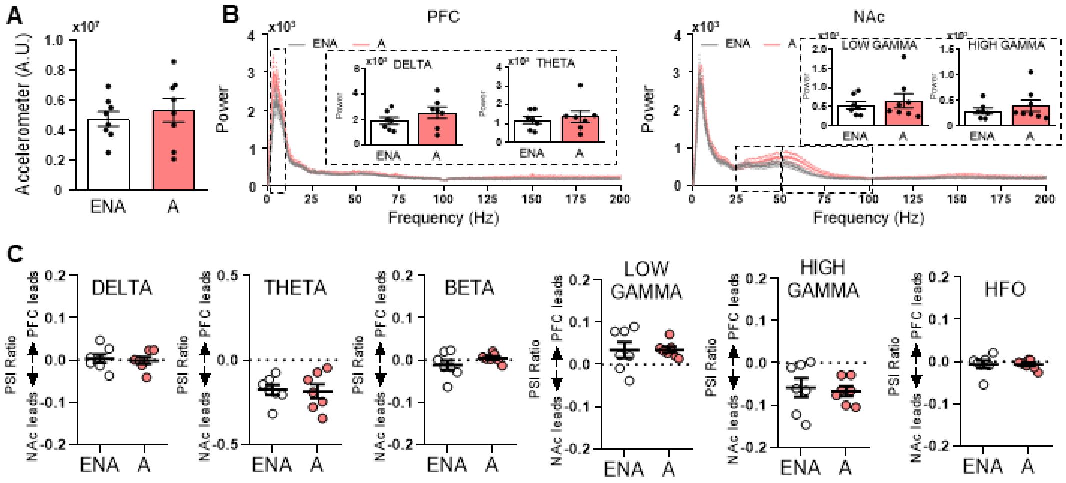


**Supplementary Fig. 2 Addicted and exteme non-addicted mice exhibited a flow of information at theta, high gamma,and low gamma frequencies under basal conditions. A** Locomotor activity in extreme non-addicted (ENA) and addicted (A) mice during recordings in the standard cage after FR5 sessions. Accelerometer, variance of signals from the accelerometer integrated within the headstages. The corresponding quantification of the animal's mobility is presented as a ratio to the highest value in the baseline condition. **B** Power spectra of PFC and NAc signals during recordings of 20 min in the standard cage. Power values of delta and theta between ENA and A mice are shown in PFC power spectra. Also, power values of lgamma and hgamma between ENA and A mice are shown in NAc power spectra. **C** PFC-NAc circuit communication (PSI) at delta, theta, beta, lgamma, hgamma and hfo bands in ENA and A animals during recordings in the standard cage.

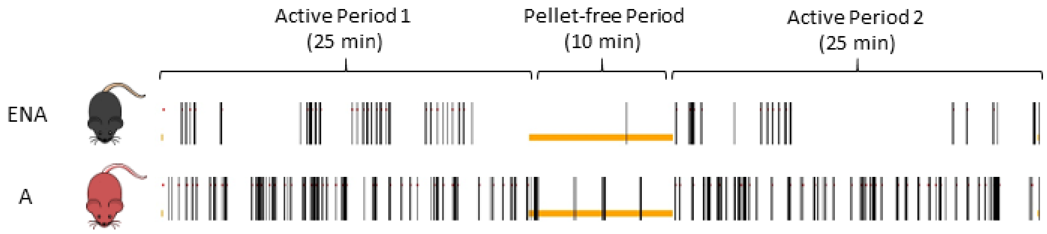


**Supplementary Fig. 3. Addicted mice performed a higher number of lever-presses compared to extreme non-addicted mice during FR5 post-surgery sessions.** Representative raster plot of the common FR5 session. Black colored vertical lines indicate lever-presses. Red dots indicate the delivery of rewards. The orange bar represents the illumination period of the operant box during the pellet-free period.

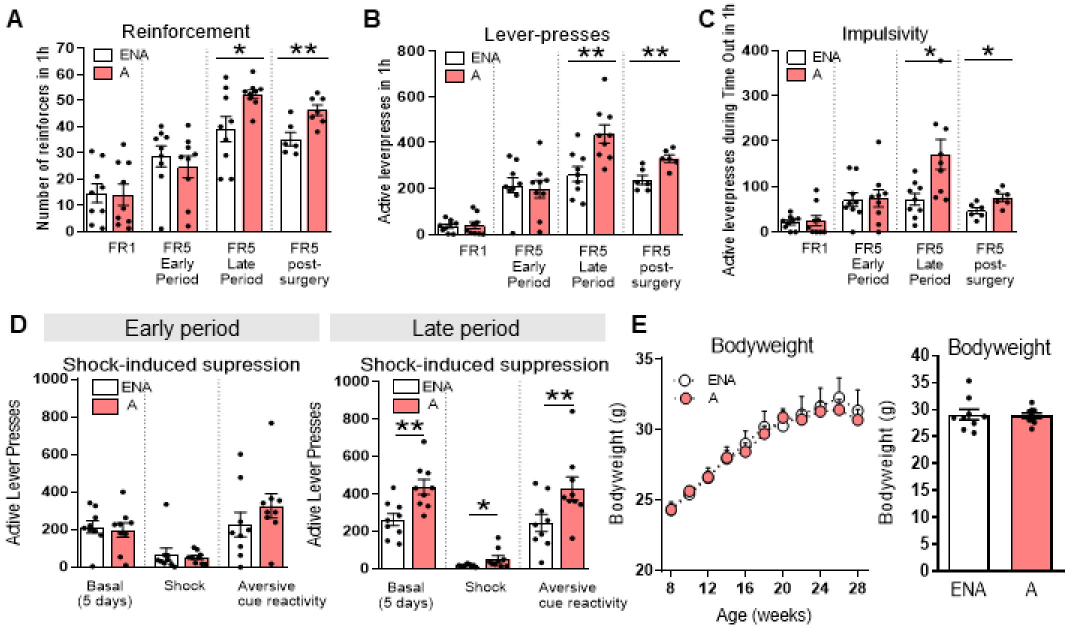


**Supplementary Fig. 4 A-C Addicted mice maintained the addictive-like behavior after the surgery period. A** number of reinforcers, **B** number of lever-presses and **C** impulsivity. Behavioral data of phenotypic traits in FR5 sessions are represented by individual values with the mean and ±SEM. FR1 corresponds to the five operant sessions of FR1, FR5 Early Period corresponds to the five FR5 sessions before the early period, FR5 Late Period corresponds to the five FR5 sessions before the late period. FR5 post-surgery corresponds to the ten FR5 recording sessions after electrode implantation, recovery and habituation. **A** Reinforcement levels quantified by counting the number of reinforcers consumed in 1h operant sessions. **B** Number of total active lever-presses scored in 1h operant session. **C** Impulsivity. Number of non-reinforced active lever-presses during five consecutive daily time out (15 s) after each pellet delivery (unpaired t-test, *P < 0.05, **P < 0.01). **D** Considering the phenotypic categorization used (n=9 A, n=9 ENA), compulsivity and aversive associative learning were re-evaluated at the early and late periods. Differences in compulsivity and aversive associative learning were observed at the late period, being A mice more compulsive, showing an increased aversive associative learning compared with ENA mice. Behavioral test of shock-induced suppression at the early period and late period represented by individual values with the mean of active levers and ±SEM. Basal corresponds to the five FR5 sessions before the compulsivity test. Shock corresponds to the compulsivity test where each pellet delivery was associated with a punishment, an electric foot shock. Shock-associated cue corresponds to the number of non-reinforced active responses in 50 min in the following session after the shock test with the same discriminative stimulus (grid floor) as shock test in which pressing the active lever had no consequences: no shock, no reward delivery and no cue light (unpaired t-test, *$P < 0.05$, **$P < 0.01$). **E** Bodyweight. Previous results were not affected by the body weight variable, as no differences in body weight were observed between the extreme subgroups. Measurements of body weight in grams across weeks in ENA and A mice (left). Averaged body weight data per phenotype is also shown (right).

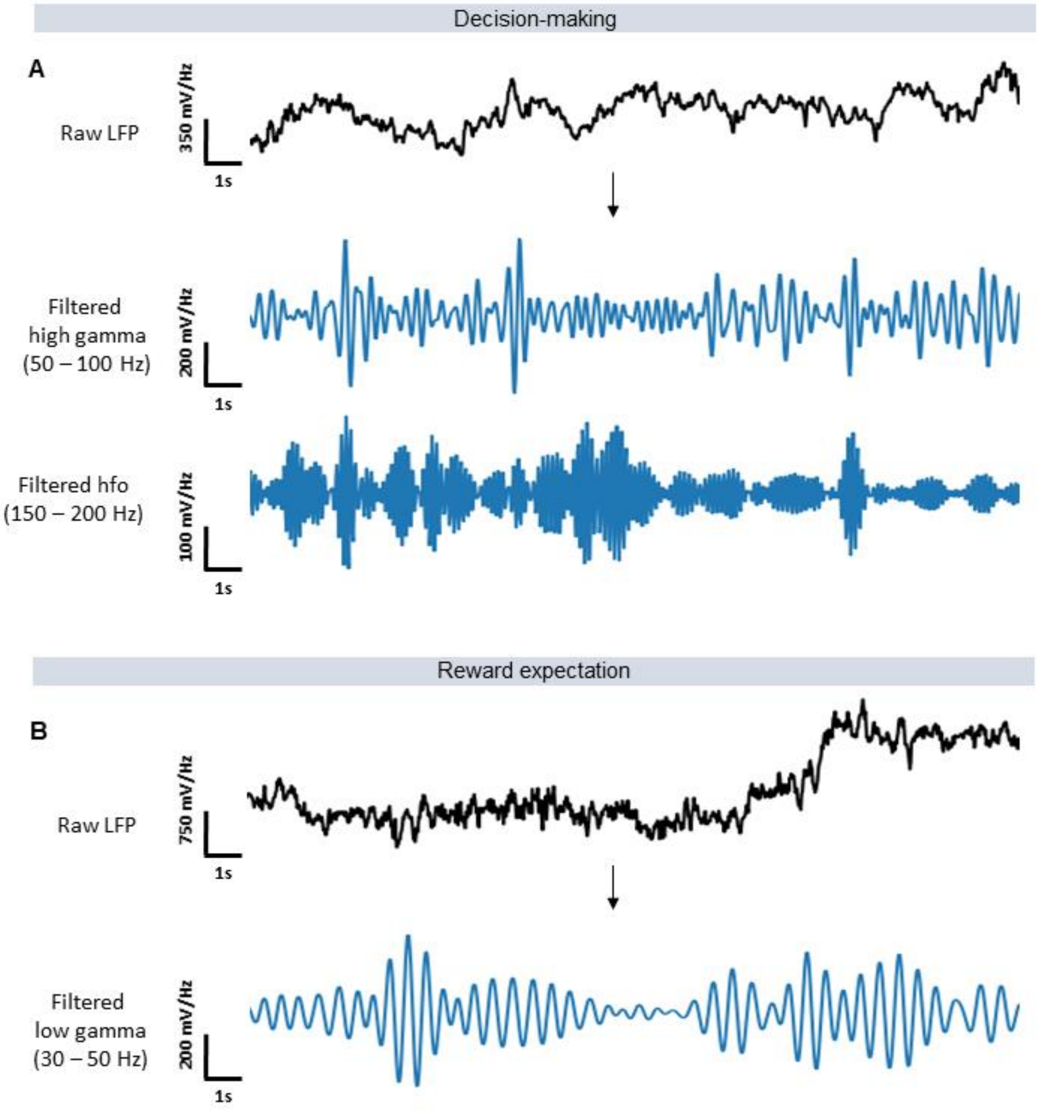


**Supplementary Figure 5. Neural signals recorded during decision-making and reward expectation. A** Representative example of raw local field potential (LFP) during decision-making (1s), filtered high gamma (50-100 Hz) and hfo (150-200 Hz) components of the signal. **B** Representative example of raw LFP during reward expectation (1s), filtered low gamma (30-50 Hz) components of the signal.

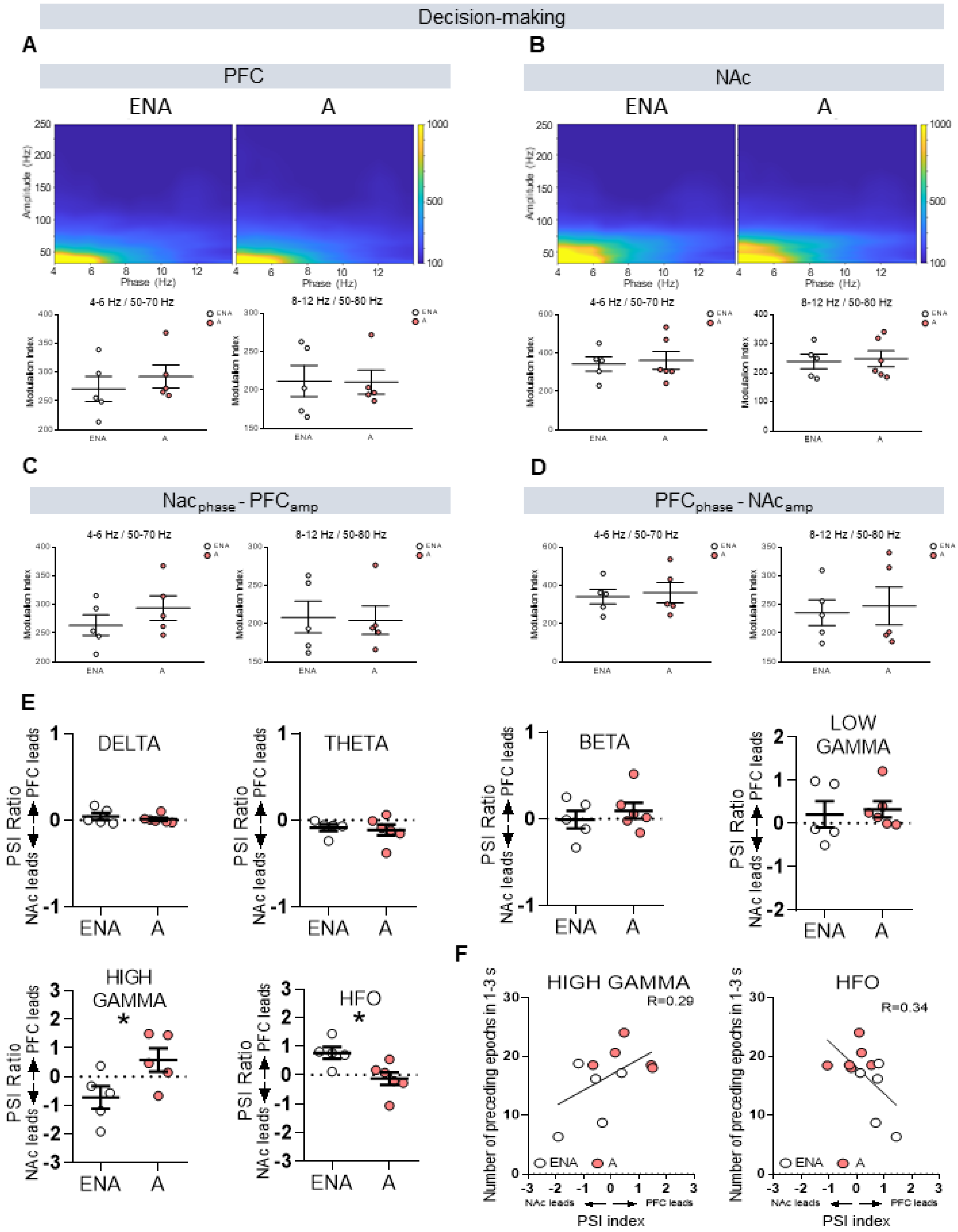


**Supplementary Fig. 6 Prefrontal-accumbens synchrony at gamma and high frequencies are disrupted in addicted mice during decision-making. A-D** Local and inter-regional cross-frequency coupling during decision-making (1s) in ENA and A mice. **A-B** Comodulation maps and quantification of delta-gamma and theta-gamma local coupling in the PFC and the NAc. The x-axis represents phase frequencies (4-14 Hz) and the y-axis represents amplitude frequencies (30-250 Hz). **C-D** Quantification of inter-regional modulation index between the NAc phase (NAcp) and the PFC amplitude (PFCa) and between the PFC phase (PFCp) and the NAc amplitude (NAca). **E** PFC-NAc circuit communication (PSI) in ENA and A animals during decision-making (unpaired t-test, $*P < 0.05$). **F** Pearson correlations between PFC-NAc circuit communication at hgamma and hfo; and the number of 1-3s preceding epochs in the FR5 operant sessions. Data are represented by individual values with the mean ±SEM.

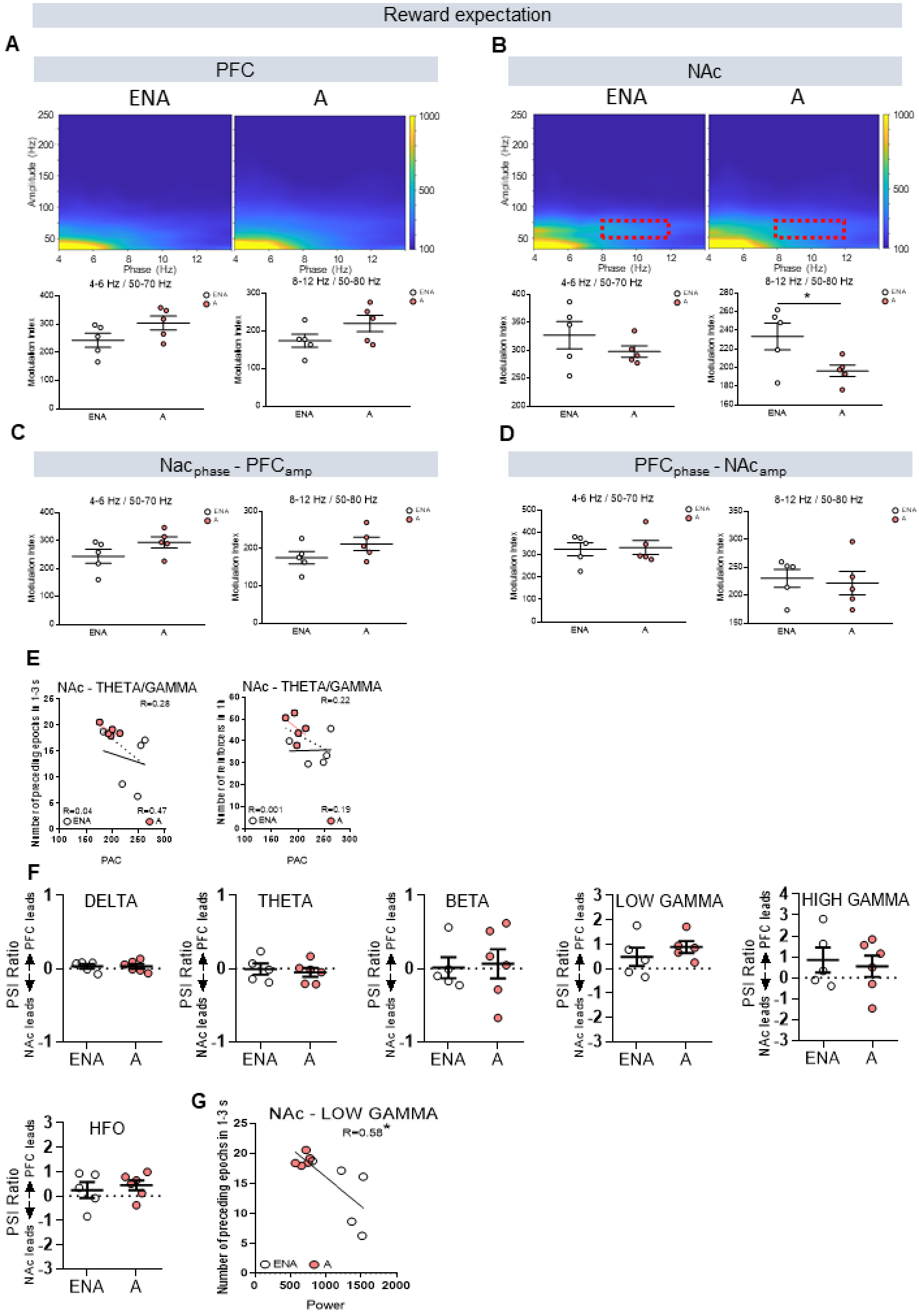


**Supplementary Fig. 7 Prefrontal-accumbens synchrony at gamma and high frequencies are disrupted in addicted mice during reward expectation. A-D** Local and inter-regional cross-frequency coupling during reward expectation (1s) in ENA and

A mice. **A-B** Comodulation maps and quantification of delta-gamma and theta-gamma local coupling in the PFC and the NAc (unpaired t-test, *P < 0.05). The x-axis represents phase frequencies (4-14 Hz) and the y-axis represents amplitude frequencies (30-250 Hz). **C-D** Quantification of inter-regional modulation index between the NAc phase (NAcp) and the PFC amplitude (PFCa) and between the PFC phase (PFCp) and the NAc amplitude (NAca). **E** Pearson correlations between local NAc theta-gamma coupling and the number of 1-3s preceding epochs, and between reinforcement levels acquired in the FR5 post-surgery operant sessions. **F** PFC-NAc circuit communication (PSI) in ENA and A animals during reward expectation. **G** Pearson correlations between NAc lgamma power and the number of 1-3s preceding epochs acquired in the FR5 operant sessions (unpaired t-test, *P < 0.05). Lgamma power negatively correlated with the number of selected preceding epochs in ENA and A mice together.

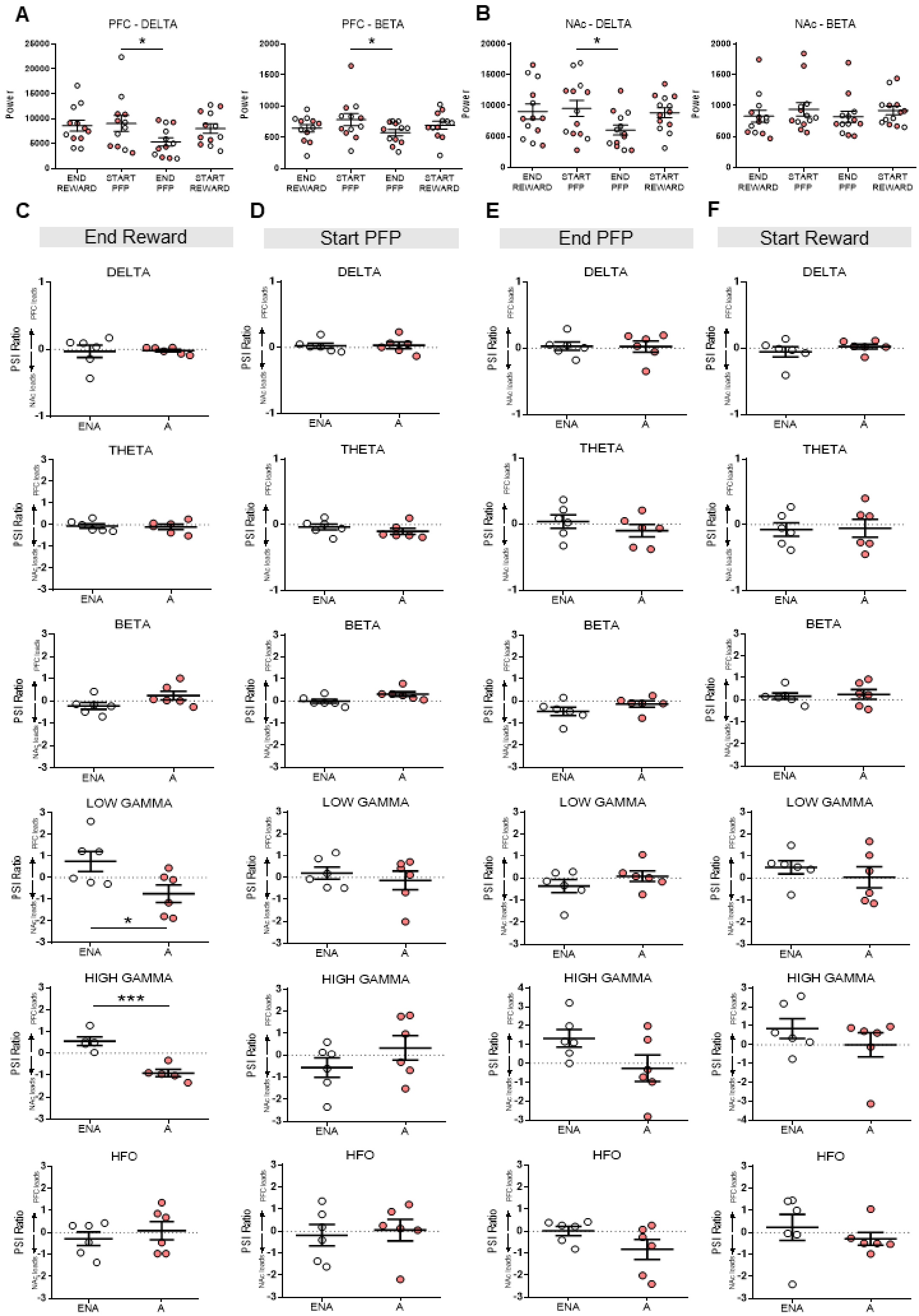


**Supplementary Fig. 8 Addicted mice exhibit abnormal prefrontal-accumbens neural dynamics during active rewarding periods. A-B** Power quantification during the phases of the operant session at delta and beta in the **A** PFC and the **B** NAc. ENA mice correspond to white colored dots whereas A mice are represented with red colored dots. Data are represented as mean ±SEM (One-way ANOVA repeated measures, *P < 0.05). **C-F** PFC to NAc circuit communication (PSI) in ENA and A mice during the end of the **c** rewarding period (end reward) (unpaired t-test, *P < 0.05, ***P < 0.001) ,the **D** start of the pellet-free period, the **E** end of the pellet-free period, and the **F** beginning of the second active period (AP2) (start reward). Differences are reported as mean ±SEM.

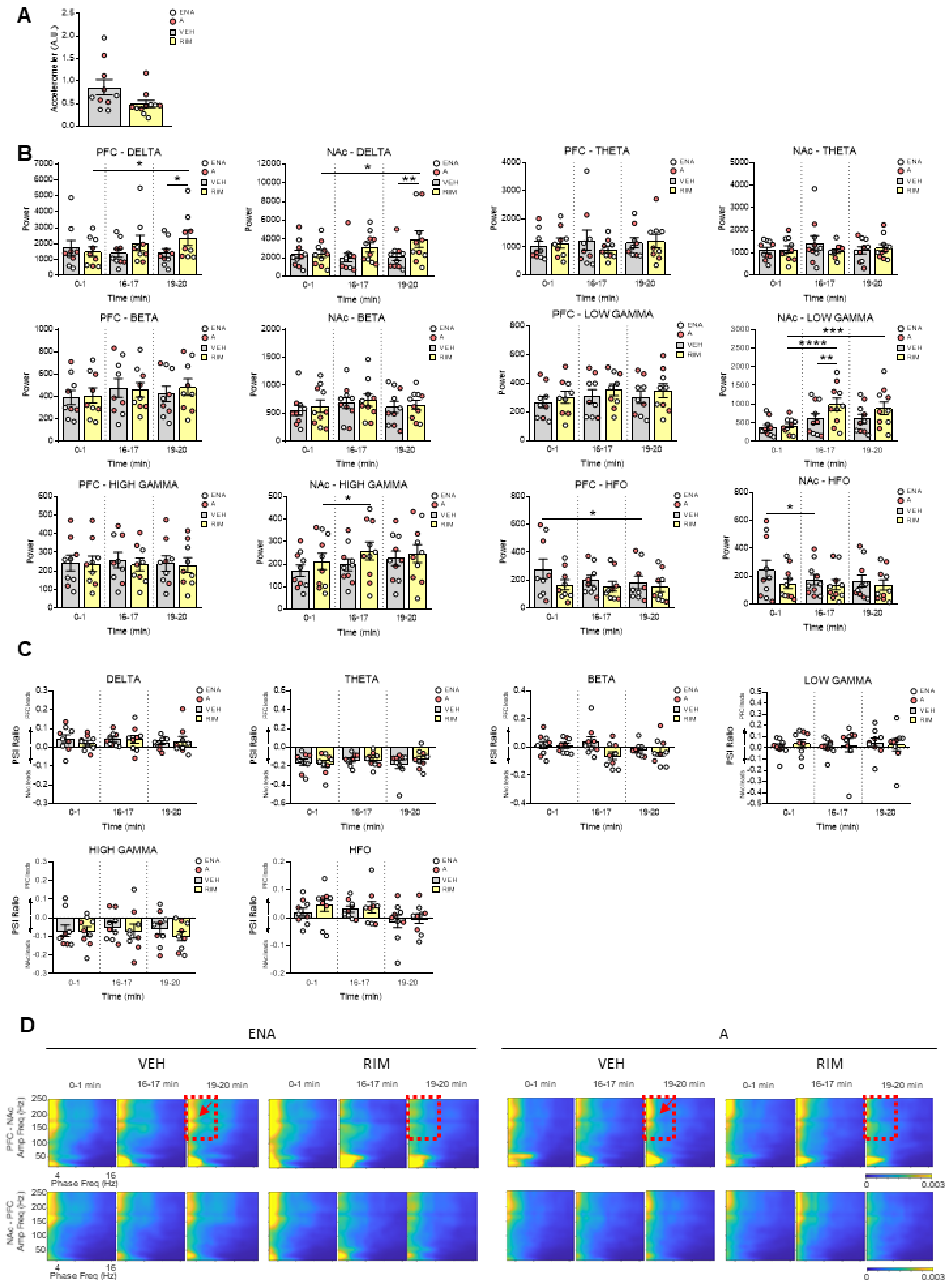


**Supplementary Fig. 9 CB1 receptors shape prefrontal–accumbens neural dynamics.** **A** Locomotor activity in ENA (white dots) and A (red dots) mice during 20 min recordings in the standard cage after vehicle (grey) or rimonabant (yellow) administration. Accelerometer, variance of signals from the accelerometer integrated within the headstages. The corresponding quantification of the animal's mobility is presented as a ratio to the highest value in the baseline condition. **B** Power quantification across time (0-1, 16-17 and 19-20 min) after administration of vehicle or rimonabant in the PFC and the NAc (two-way ANOVA multiple comparisons, *P < 0.05). **C** PFC-NAc circuit communication (PSI) in ENA and A mice across time (0-1, 16-17 and 19-20 min after administration of vehicle or rimonabant). Data is reported as mean ±SEM. **D** Comodulation maps quantifying inter-regional modulation index between the PFC phase (PFCp) and the NAc amplitude (NAca) and between the NAc phase (NAcp) and the PFC amplitude (PFCa) cross-frequency coupling in 1 minute periods (0-1, 16-17 and 19-20 min) in ENA and A mice. The x-axis represents phase frequencies (2-16 Hz) and the y-axis represents amplitude frequencies (10-250 Hz). Numbers on top indicate the minute after vehicle or rimonabant administration. The red dotted square indicates the coupling between delta and hfo present in normal conditions (vehicle), marked with a red arrow, that disappear after 19 minutes of rimonabant administration.

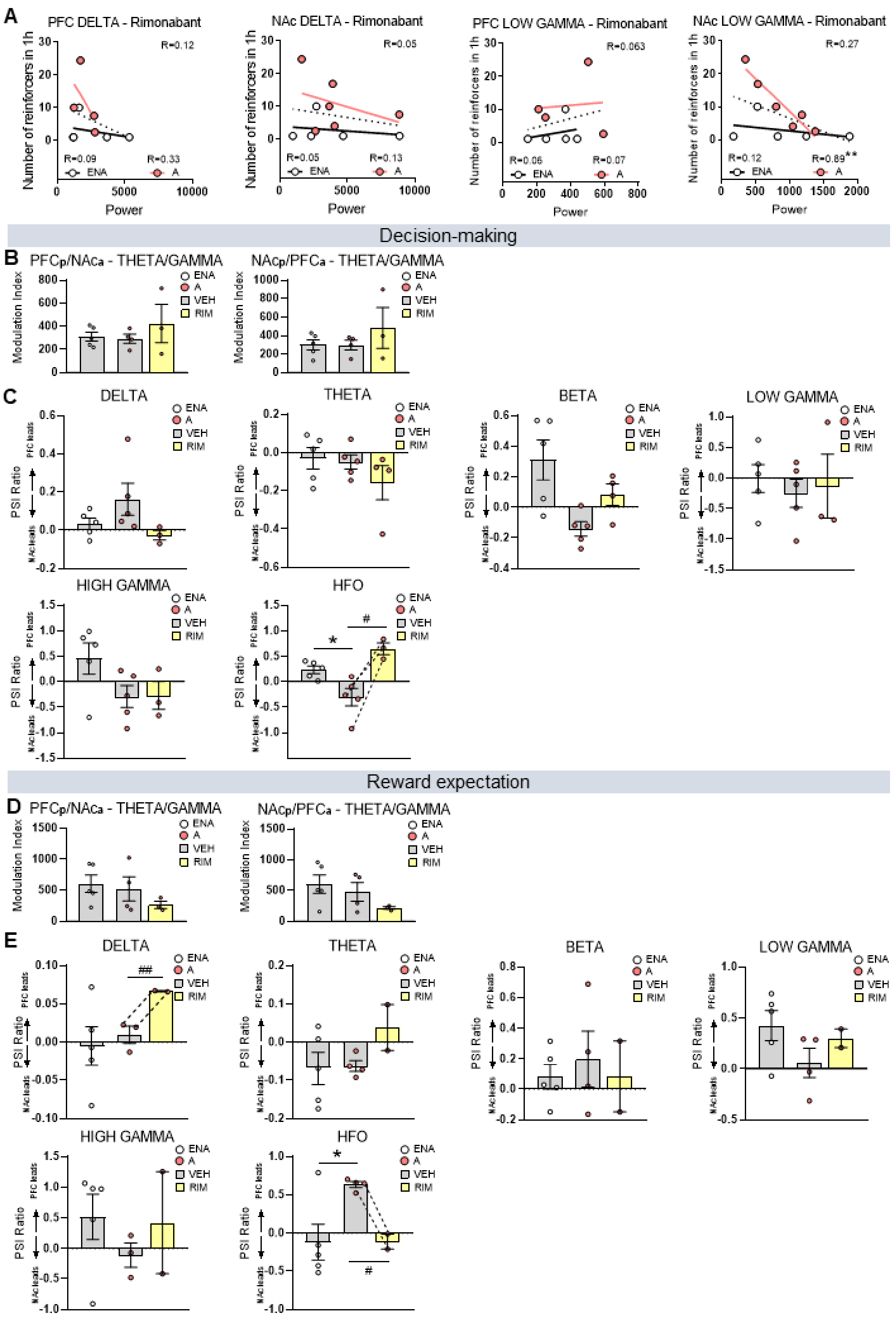


**Supplementary Fig. 10 Rimonabant reduces reinforcement levels and reverses abnormal prefrontal-accumbens neural dynamics during decision-making and reward expectation. A** Pearson correlations between the increase of delta and lgamma power observed after 19 minutes of rimonabant administration and reinforcement levels of the subsequent FR5 post-surgery operant sessions. The dotted line corresponds with linear regression of extreme non-addicted (ENA) and addicted (A) mice together, whereas linear regressions in ENA and A mice separately are represented in black and red colored lines, respectively. **B** Quantification of theta-gamma inter-regional coupling during decision-making (1s) in ENA and A mice treated with vehicle and rimonabant treated A mice. **C** PFC-NAc circuit communication (PSI) during decision-making in ENA and A mice treated with vehicle and rimonabant treated A mice (unpaired and paired t-test, *P $< 0.05$, #P $< 0.05$). **D** Quantification of theta-gamma inter-regional coupling during reward expectation (1s) in ENA and A mice treated with vehicle and rimonabant treated A mice. **E** PFC-NAc circuit communication (PSI) during reward expectation in ENA and A mice treated with vehicle and rimonabant treated A mice.
