## Supplementary figures and images for "Prefrontal–accumbens neural dynamics abnormalities in mice vulnerable to develop food addiction"

### SI Figures

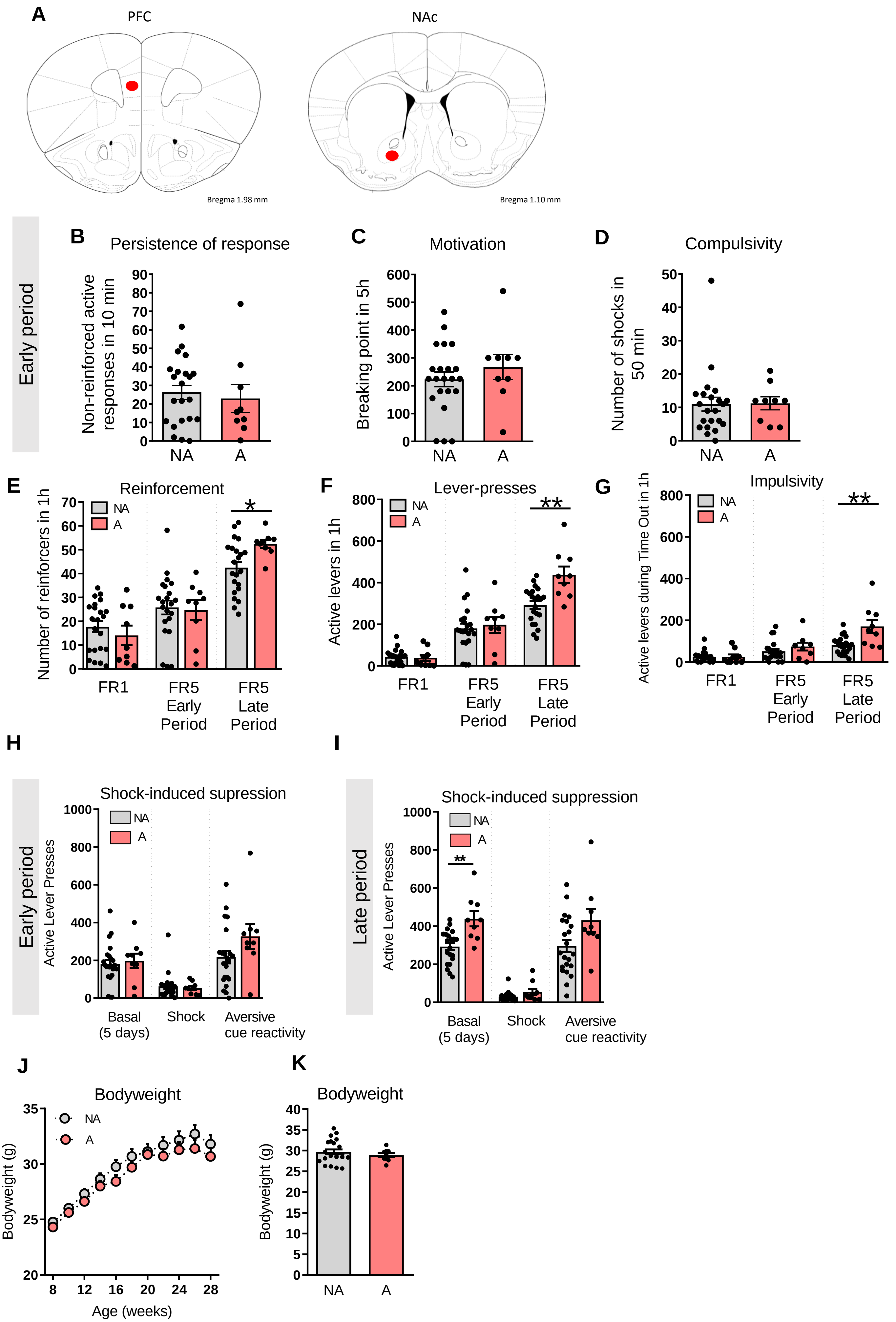

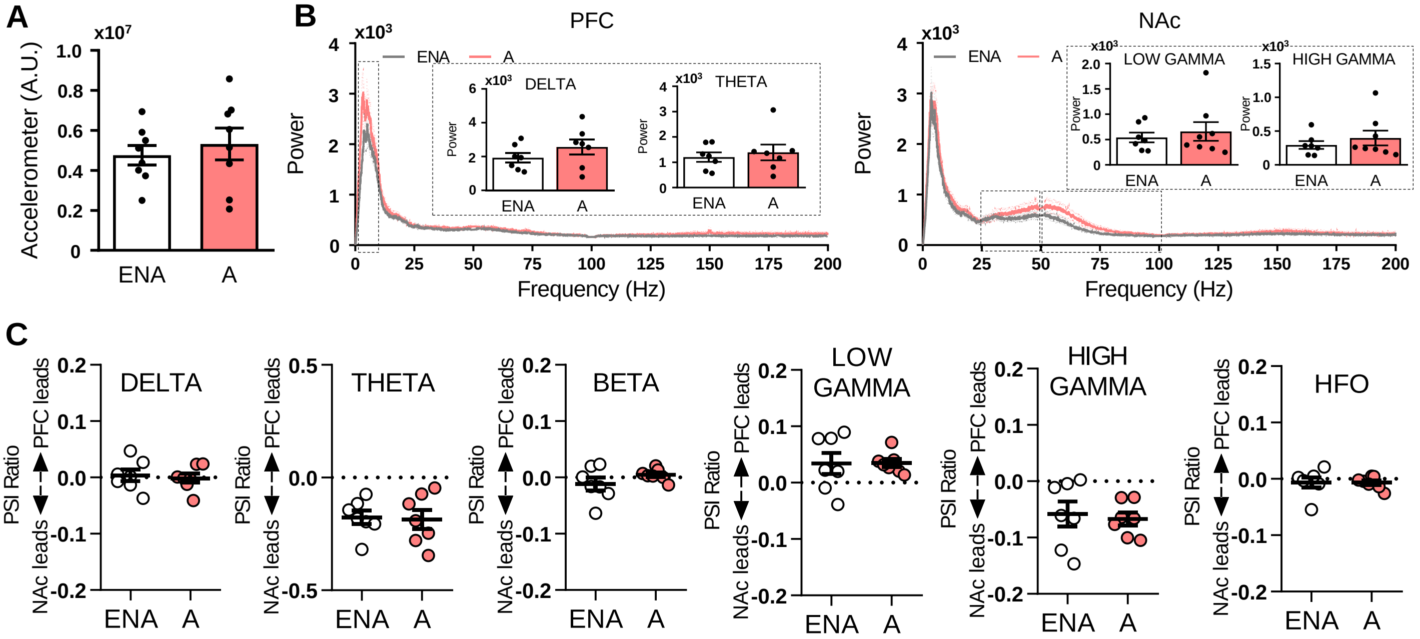

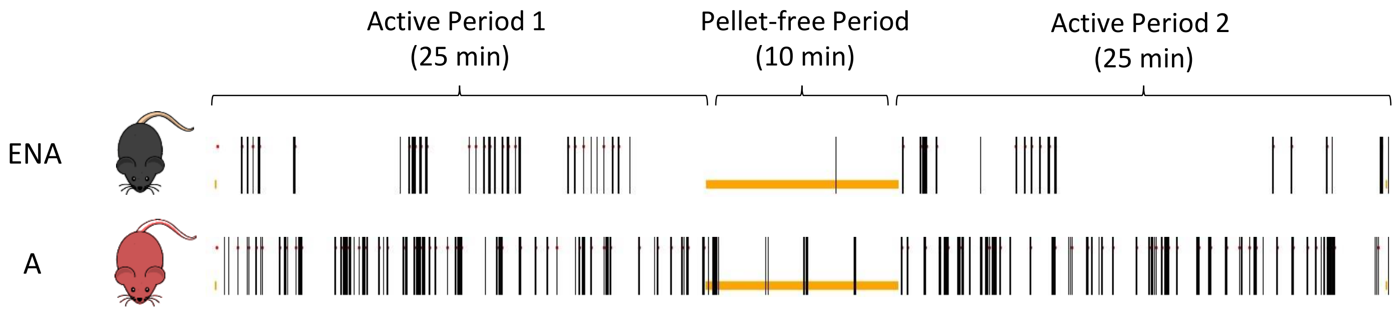

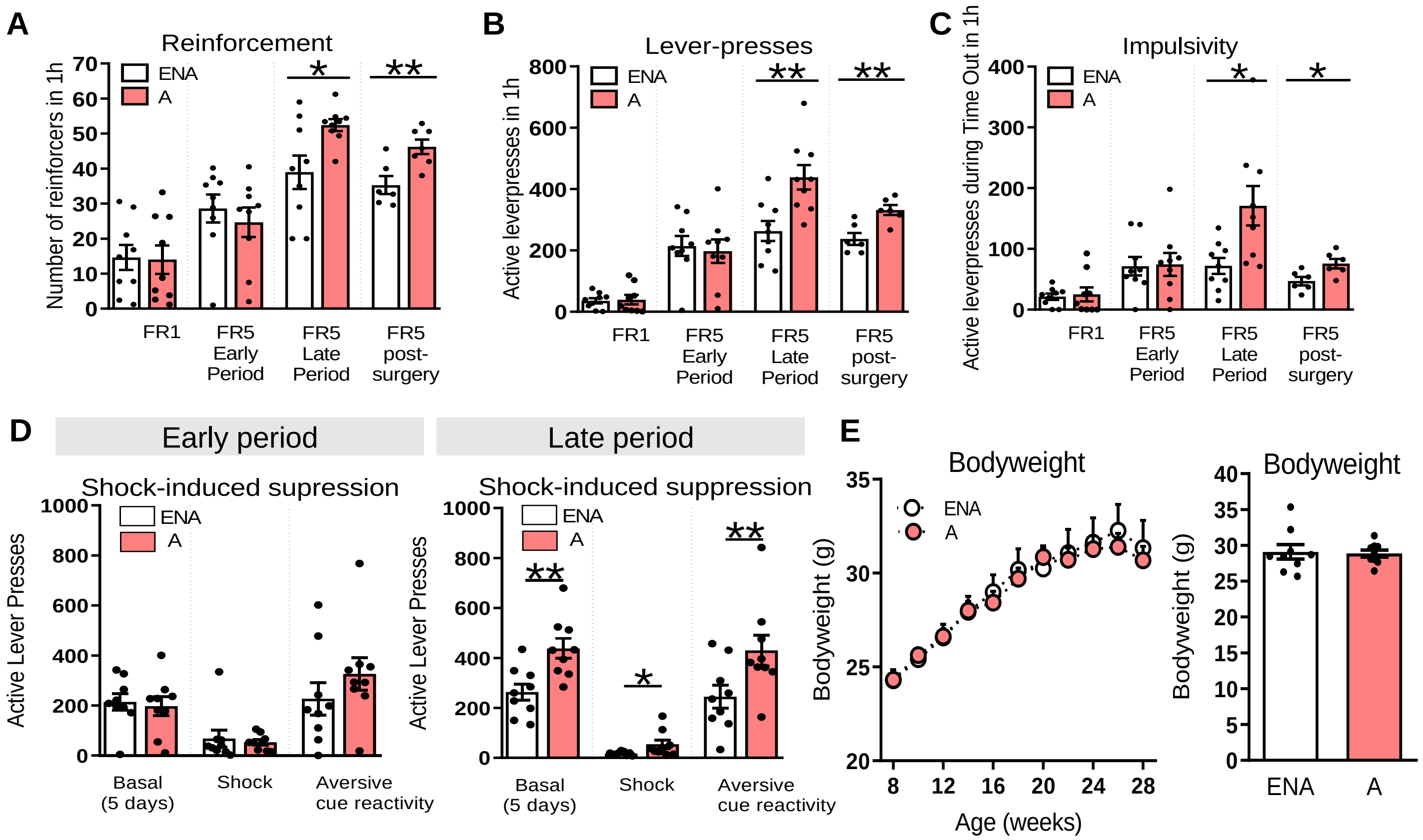

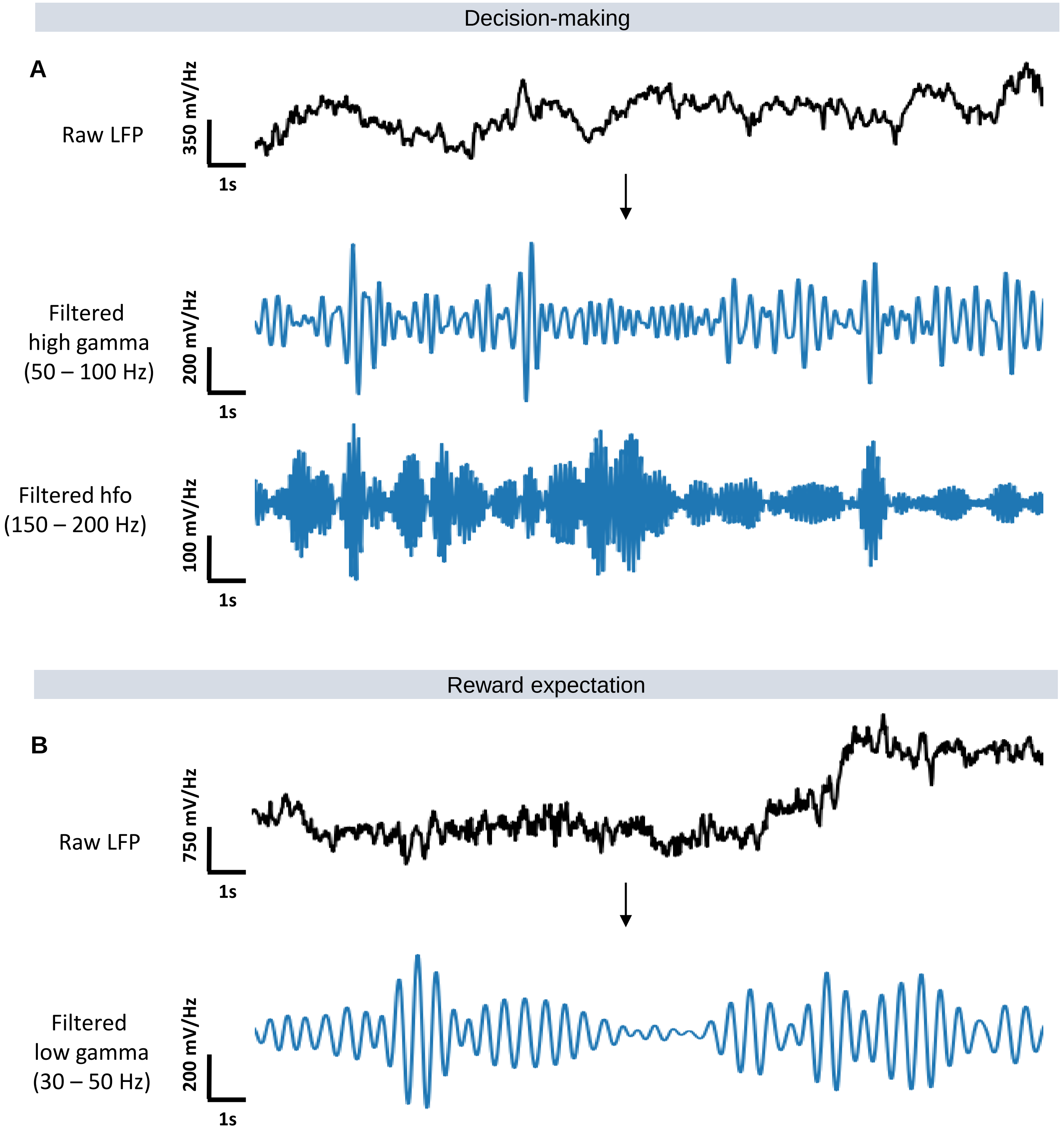

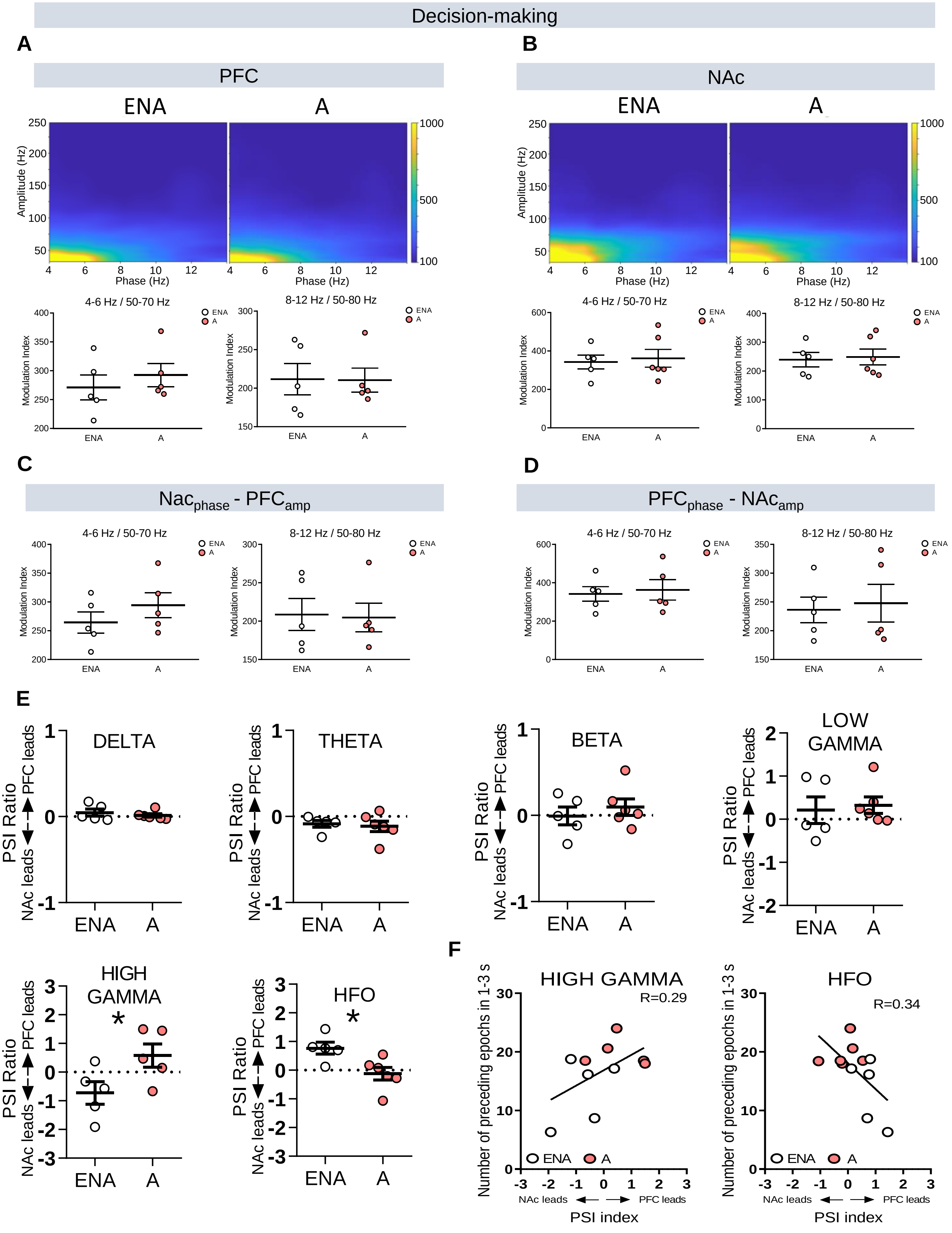

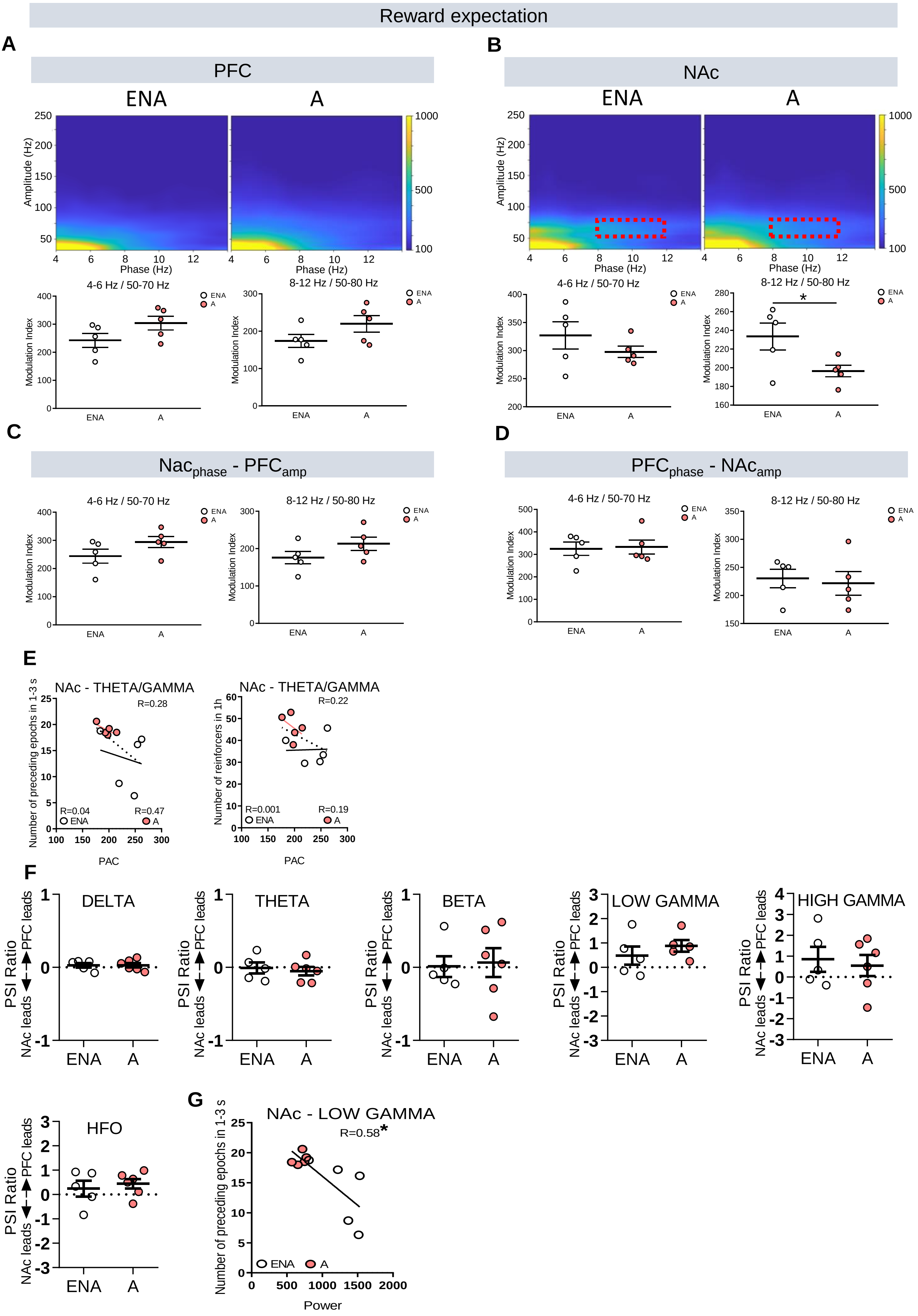

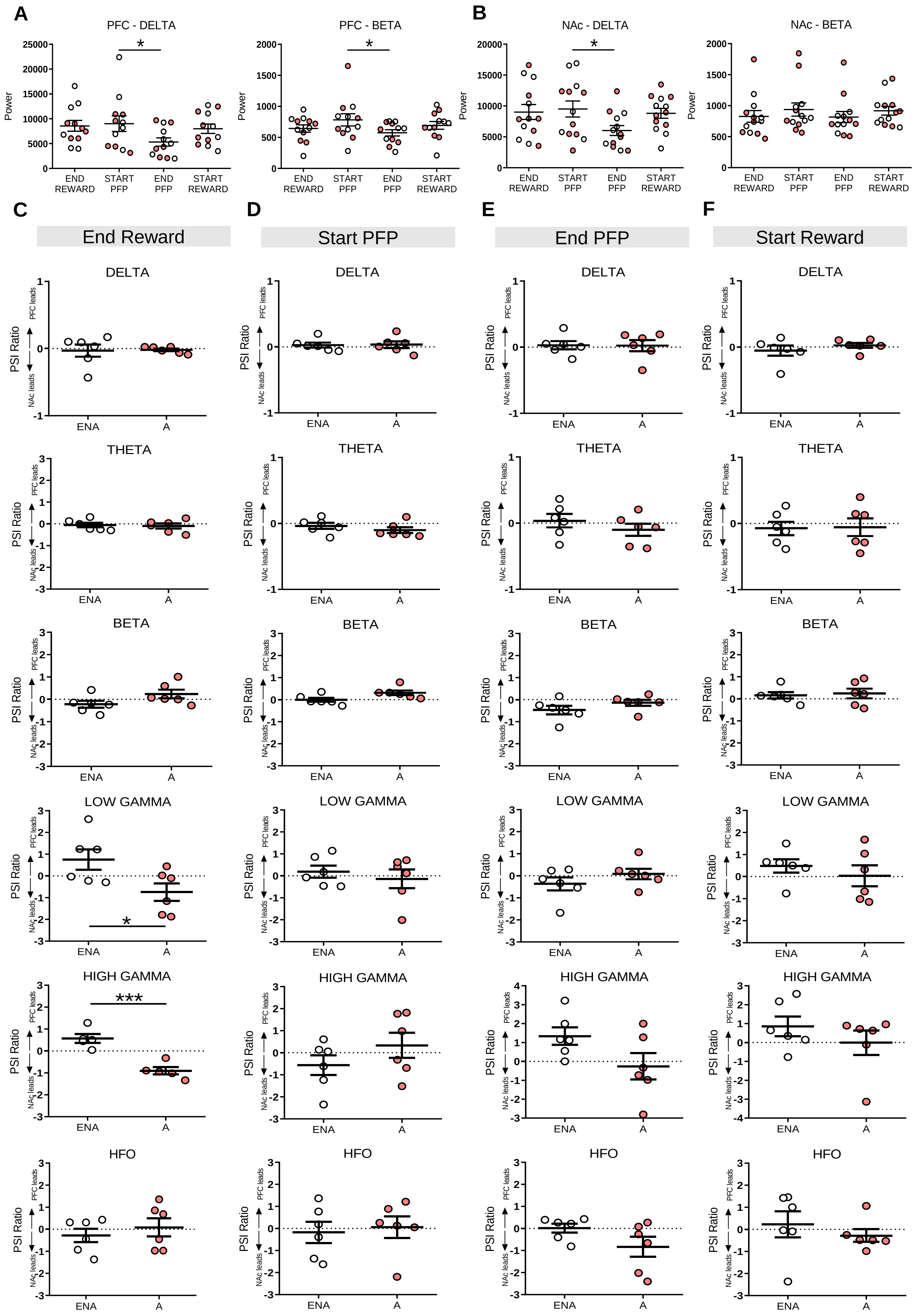

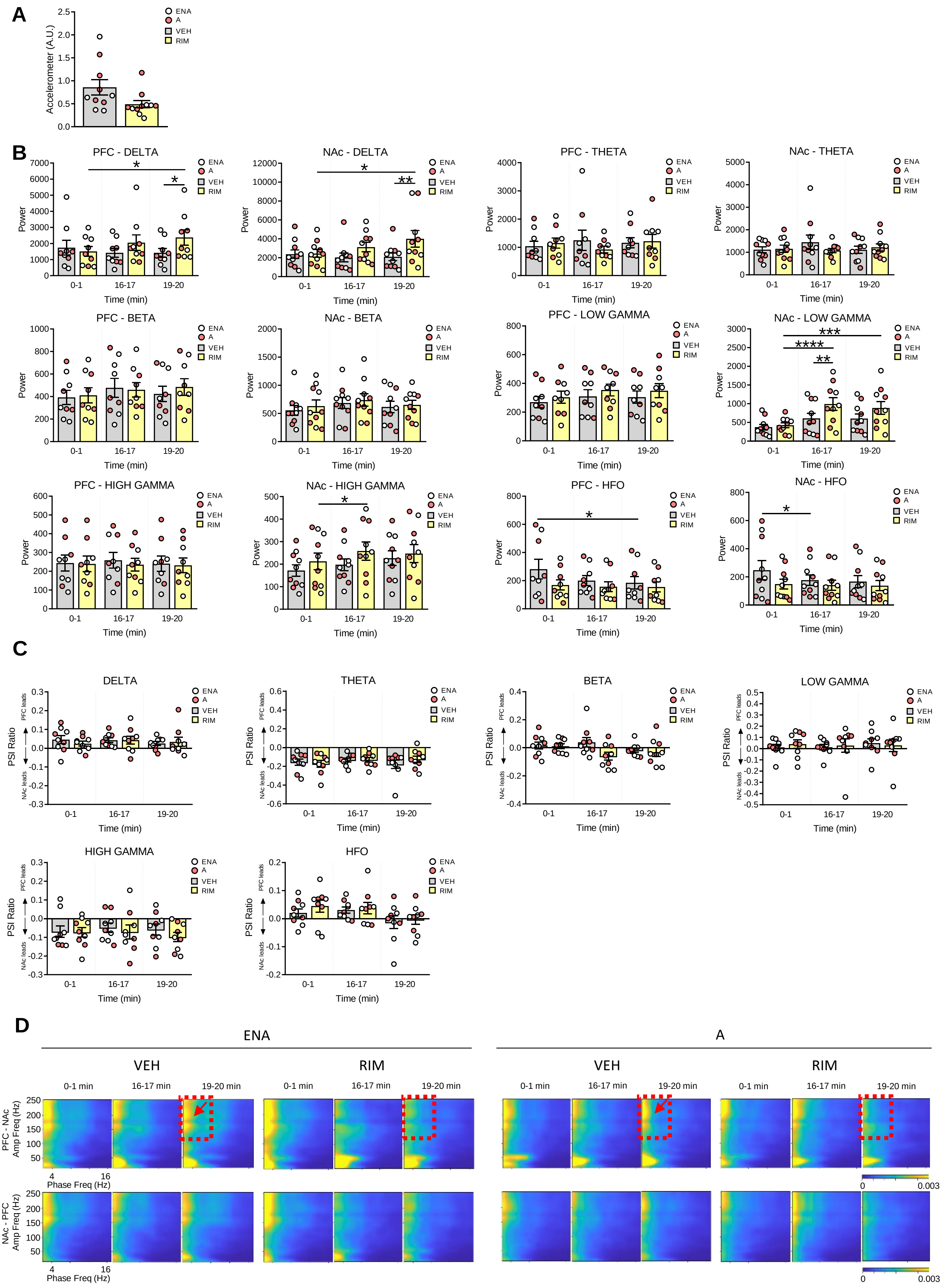

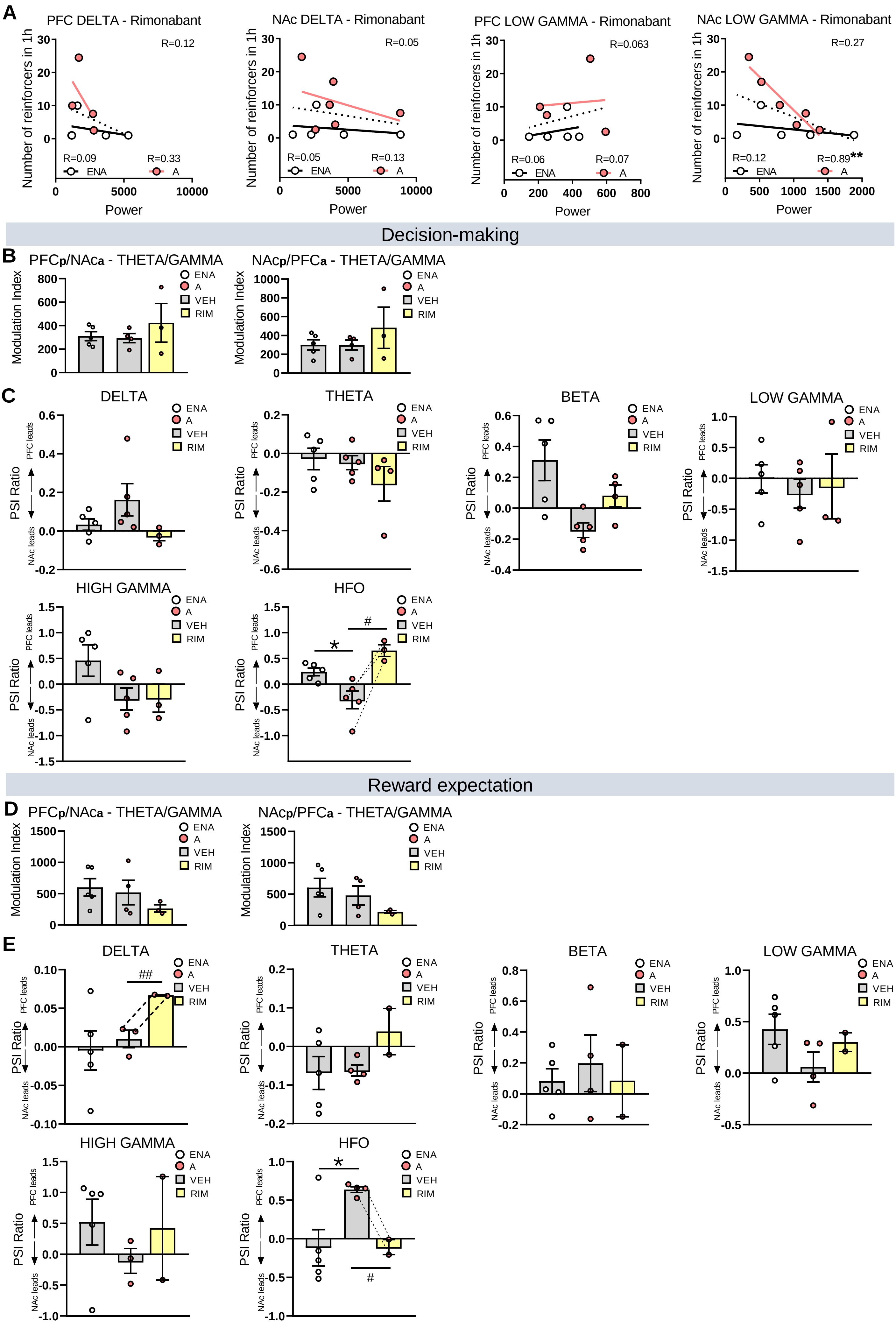
